# The Neural System of Metacognition Accompanying Decision-Making in The Prefrontal Cortex

**DOI:** 10.1101/135772

**Authors:** Lirong Qiu, Jie Su, Yinmei Ni, Yang Bai, Xiaoli Li, Xiaohong Wan

**Affiliations:** State Key Laboratory of Cognitive Neuroscience and Learning and IDG/McGovern Institute for Brain Research, Beijing Normal University, Beijing, 100875, China.

**Keywords:** Metacognition, Decision-making, Uncertainty, Cognitive control, Dorsal anterior cingulate cortex, Frontopolar cortex, Prefrontal cortex, fMRI

## Abstract

Decision-making is usually accompanied by metacognition, through which a decision maker monitors the decision uncertainty and consequently revises the decision, even prior to feedback. However, the neural mechanisms of metacognition remain controversial: one theory proposes that metacognition coincides the decision-making process; and another addresses that it entails an independent neural system in the prefrontal cortex (PFC). Here we devised a novel paradigm of “decision-redecision” to investigate the metacognition process in redecision, in comparison with the decision process. We here found that the anterior PFC, including dorsal anterior cingulate cortex (dACC) and lateral frontopolar cortex (lFPC), were exclusively activated after the initial decisions. dACC was involved in decision uncertainty monitoring, whereas lFPC was involved in decision adjustment controlling, subject to control demands of the tasks. Our findings support that the PFC is essentially involved in metacognition and further suggest that functions of the PFC in metacognition are dissociable.

## Introduction

Decision-making is a process of evidence accumulation. The evidence comes from sensory signals of external stimuli or mental representations of internal cognitive operation. Variations of evidence may render a decision uncertain. A decision maker is often intentionally or automatically aware of such an uncertain state of the decision, and confirms or revises the initial decision, even prior to feedback. For instance, before submitting the manuscript, the authors have revised it several times, as being aware of uncertainty, although the review outcome is unknown. In literature, the processes of decision uncertainty monitoring and consequent decision adjustment are termed as metacognition, that is, “cognition about cognition” (Flavell, 1979; Nelson and Narens, 1990; Dunlosky and Metcalfe, 2009; Fleming and Dolan 2012). Although metacognition usually accompanies decision-making, the underlying neural processes of decision uncertainty monitoring and consequent decision adjustment remain less clear than that of the decision process per se (Gold and Shadlen, 2008; Rushworth et al., 2011), and might be misattributed to the decision-making process.

Much of the work on neural basis of metacognition has focused on metacognitive monitoring of internal states (i.e., confidence, or uncertainty) of such cognitive processes as episodic memory (Kikyo et al., 2002; Chua et al., 2006) and sensory perception in human (Fleming et al., 2010, 2012b; Resulaj et al., 2009; Kiani et al., 2014; van den Berg et al., 2016; Murphy et al., 2016), as well as sensory perception in animals (Kepecs et al., 2008; Kiani and Shadlen, 2009; Middlebrooks and Sommer, 2012; Komura et al., 2013). Behaviorally, the confidence ratings that reflect subjective accuracy beliefs on decisions were often found to deviate from the actual decision accuracy (Kunimoto et al., 2001; Lau and Passingham, 2006; Wilimzig et al., 2008; Song et al., 2011). These observations indicate that there should exist a separate neural system (meta-level) to monitor the decision process (object-level) (Flavell, 1979; Nelson and Narens, 1990; Dunlosky and Metcalfe, 2009; Fleming and Dolan, 2012). The prefrontal cortex (PFC) has been suggested to play critical roles in the metacognitive monitoring of decisions (Kikyo et al., 2002; Chua et al., 2006; Shimamura, 2008; Del Cul et al., 2009; Rounis et al., 2010; Fleming et al., 2010, 2012; Ham et al., 2014; Wan et al., 2016). Essentially, interference or lesions of the PFC merely impaired the ability of metacognitive monitoring of decisions, but not the decisions per se (Del Cul et al., 2009; Rounis et al., 2010; Ham et al., 2014; Fleming et al., 2014).

On the contrary, it has been addressed that metacognition could be merely dependent on the decision-making process, and exclusively relies on accumulated evidence (Vickers 1979; Kiani and Shadlen, 2009; Resulaj et al., 2009; Pleskac and Busemeyer, 2010; Kiani et al., 2014; Yu et al., 2015; van den Berg et al., 2016). Specifically, this theory on the basis of bounded accumulation models interpreted that the divergence between decision accuracy and confidence reports might be caused by continuous post-decisional evidence accumulation during the intervals between decisions and confidence reports (Resulaj et al., 2009; Pleskac and Busemeyer, 2010; Yu et al., 2015; van den Berg et al., 2016). Further, decision adjustment could naturally occur by continuous post-decisional evidence accumulation (Resulaj et al., 2009; van den Berg et al., 2016). Therefore, it argues that a separate neural system for metacognition to monitor and control the decision-making process should be not necessary (van den Berg et al., 2016).

The purpose of the retrospective metacognition accompanying uncertain decisions is to confirm or revise the foregone decisions, prior to feedback. Given an opportunity to make a decision on the same situation again (*redecision*), the decision maker might revise the initial decision and update the confidence rating, on the basis of the foregone decisions (van den Berg et al., 2016; Wan et al., 2016). It could be difficult to discriminate the above two theories in a single decision paradigm, as the decision-making process and the metacognition process are inevitably coupled together. To examine the behavioral performance and neural activities in redecision, however, may allow us to directly test whether the metacognition process would coincide the decision-making process or entail another separate neural system (Yeung and Summerfield, 2014; Fleming, 2016). If it were the former, then the redecision process would evoke exactly the same neural system of the decision-making process as that in the initial decision. Critically, the divergence between decision accuracy and confidence reports within and across individual participants would be much reduced, or the individual metacognitive abilities would be much improved by redecision, as more evidence would be further accumulated. Otherwise, a separate neural system for metacognition, other than that occurred in the initial decision, would be newly recruited in redecision. Importantly, the individual metacognitive abilities might be intrinsically dependent on the circuit of this separate neural system, other than that of the decision-making system or the accumulated evidence. In other words, the individual metacognitive abilities would be not much changed by redecision. Further, in addition to metacognitive monitoring that immediately occurs after decisions even with no requirement for redecision (Wan et al., 2016), the process of redecision should be necessarily comprised of metacognitive controlling, to revise the foregone decisions, which should be different from the decision-making process in the initial decision. Metacognitive monitoring and metacognitive controlling are the two key components of metacognition (Flavell, 1979; Nelson and Narens, 1990; Dunlosky and Metcalfe, 2009). So far, the neural process of metacognitive controlling has been little explored (Wan et al., 2016).

In the present study, we employed a novel experimental paradigm – “decision-redecision” (Figure 1A). The participants made two consecutive decisions on the same situation in a perceptual decision-making task and a rule-based decision-making task. We employed this new paradigm in functional magnetic resonance imaging (fMRI), to systematically investigate the neural processes of metacognition during the redecision phase, in comparison with the decision-making process. We found that the metacognitive processes during the redecision phase in the two tasks commonly evoked a frontoparietal control network, including dorsal anterior cingulate cortex (dACC) and lateral frontopolar cortex (lFPC) in the anterior PFC, separate from the decision-making neural system. Critically, dACC was involved in metacognitive monitoring of decision uncertainty, encoding the subjective uncertainty states about the forgone decisions; In contrast, lFPC was involved in metacognitive controlling of decision adjustment, encoding the strategic signals for exploration of alternative options. The involvement of lFPC in metacognitive controlling was further dissociated upon the task control demand and intrinsic motivation in redecision. Thus, our findings support that the PFC is essentially involved in metacognition, which is largely independent of the decision-making neural system, and further suggest that the functions of the PFC in metacognition are dissociable.

**Figure 1.**
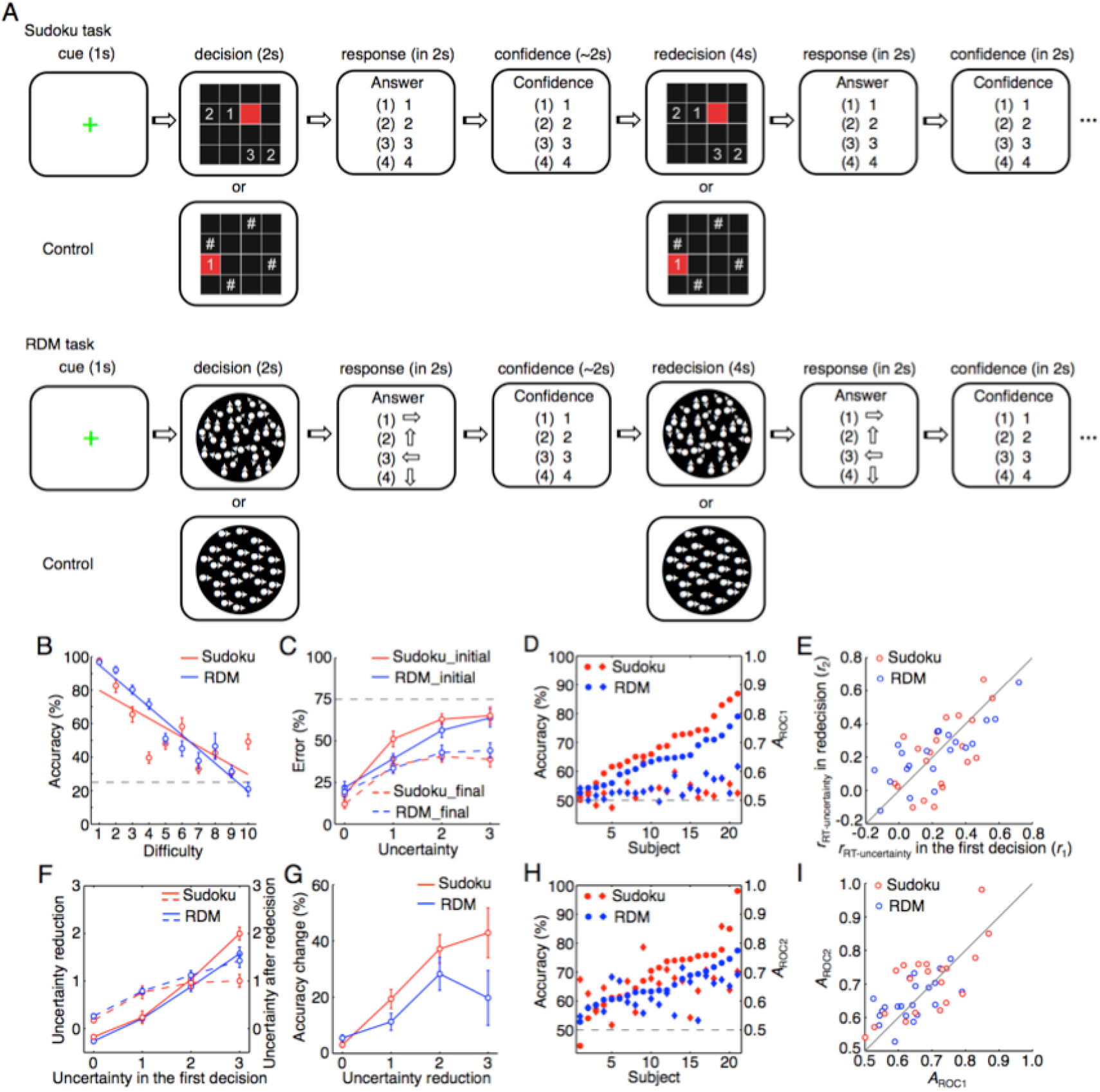
The “decision-redecision” paradigm of the Sudoku and RDM tasks and behavioral performance in the tasks. (A) The Sudoku and RDM task sequences. (B) The relationship between task difficulty and the mean accuracy in the initial decision (2-s task immediately after training). (C) The relationship between the uncertainty level of the initial decision (4 – confidence rating) and the likelihood of errors in the initial and final decisions. (D) The individual uncertainty sensitivity (*A*_ROC_, circles) and decision accuracy (diamonds) in the initial decision. (E) The individual *r*_RT-uncertainty_ in the initial and final decisions. (F) The relationship between the uncertainty level of the initial decision and the extent of uncertainty reduction by redecision (solid lines) and the uncertainty level after redecisison (broken lines). (G) The relationship between the extent of uncertainty reduction and the accuracy change by redecision. (H) The individual uncertainty sensitivity (*A*_ROC_, circles) and decision accuracy (diamonds) in the final decision. (I) The individual uncertainty sensitivity (*A*_ROC_) in the initial and final decisions. The data illustrated from C-I were from the main fMRI experiment (fMRI1). Red, the Sudoku task; Blue, the RDM task. Error bars indicate S.E.M. across the participants.

## Results

### Task paradigm

We developed a novel experimental paradigm – “decision-redecision” (Figure 1A). The participant was instructed to make an initial decision (*decision* phase), immediately followed by another decision on the same situation (*redecision* phase), so that the participant could utilize this opportunity to revise the initial decision and update the confidence rating. The internal states of uncertainty on the initial and final decisions were separately evaluated by confidence rating (four-level scales; *confidence* phase), immediately after the decisions. The uncertainty level was then negative to the confidence rating (*i.e.*, 4 – the confidence level). Different from the previous paradigm in analysis of ‘change of mind’ (Resulaj et al., 2009; van den Berg et al., 2016), which was only able to analyze a small portion of trials in which the participant happened to change the mind, our paradigm here could allow us to analyze each trial regardless of ‘change of mind.’

We used two different types of decision-making tasks in the present study; one was a rule-based decision-making (Sudoku) task, and the other was a perceptual decision-making (random dot motion, RDM) task, which had been intensively used to investigate the neural process of decision-making (Gold and Shadlen, 2008), and metacognition recently (Kiani and Shadlen, 2009; Resulaj et al., 2009; Kiani et al., 2014; van den Berg et al., 2016). The decision-making and metacognition processes of the former task might rely on internal information operation, but those of the latter might be merely dependent on accumulation of external new information. We compared the behavioral and the neural differences between the decision-making and metacognition processes, as well as their differences between the two tasks. The sequences of both tasks were identical (Figure 1A, illustrated for the main fMRI experiment, fMRI1). After a Sudoku problem or RDM stimulus was presented for 2 s, the participant made a choice from four options and then reported the confidence rating each in 2 s. Critically, the same Sudoku problem or RDM stimulus was immediately repeated for 4 s, and the participant made a choice and reported the confidence rating again each in 2 s. As the control condition, a digital number was illustrated in the target grid in the Sudoku task, and a RDM stimulus with 100% coherence was used in the RDM task. For the former, the participant only needed to press the button matching the number, and for the latter, the participant indicated the unambiguous RDM direction. For both tasks, the task difficulty (Figure 1B) of each trial was adaptively adjusted by a staircase procedure (Levitt, 1971; Fleming et al., 2010), so that the average accuracy for the first decision was converged to approximately 50% (the chance level was 25%). Prior to the experiments, each participant was trained to attain a high-level proficiency in the Sudoku problem solving.

### Behavioral results

Twenty-one participants took part in fMRI1 (See Materials and Methods). In both tasks, the uncertainty levels were largely consistent with the error likelihoods of the initial decisions (Figure 1C; *r* = 0.76 ± 0.12, mean ± standard deviation, one tailed *t* test, *t*_21_ = 7.3, *P* = 1.7 × 10^−7^ in the Sudoku task; *r* = 0.71 ± 0.14, *t*_21_ = 6.8, *P* = 5.0 × 10^−7^ in the RDM task). To examine the trial-by-trial consistency between likelihoods of erroneous decisions and the subjective belief of uncertainty in each individual participant, a nonparametric approach was employed to construct the receiver operating characteristic (ROC) curve by characterizing the error likelihoods under the different uncertainty levels of the initial decisions. The area under curve (*A*_ROC_) was calculated to represent the individual uncertainty sensitivity, indicating how precisely the participant was sensitive to the decision uncertainty (Fleming et al., 2010). As similar as the previous observations (Fleming et al., 2010; Song et al., 2011), the uncertainty sensitivities of individual participants were markedly deviated from the actual decision accuracy in both tasks (Figure 1D; one tailed paired-t test, *t*_21_ = 6.6, *P* = 7.3 × 10^−7^ in the Sudoku task; *t*_21_ = 7.8, *P* = 5.6 × 10^−8^ in the RDM task). The response time (RT) of option choices in the initial decision was strongly and positively correlated with the uncertainty level (Figure 1E; one tailed *t* test, *t*_21_ = 6.9, *P* = 4.0 × 10^−7^ in the Sudoku task; *t*_21_ = 4.3, *P* = 1.6 × 10^−4^ in the RDM task), but was weakly correlated with the task difficulty (one tailed *t* test, *t*_21_ = 2.1, *P* = 0.048 in the Sudoku task; *t*_21_ = 2.0, *P* = 0.052 in the RDM task), due to the control of task difficulties by the staircase procedure. Thus, the RT of decision here much reflected the decision uncertainty level, rather than the task difficulty, indicating that the participants should be aware of uncertainty during the choice, and might be vacillating among the options during choices. In contrast, the RT for confidence report was not correlated with the uncertainty level in both tasks (one tailed *t* test, *t*_21_ = 1.1, *P* = 0.14 in the Sudoku task; *t*_21_ = 1.2, *P* = 0.12 in the RDM task). Further, the correlation coefficient between RT of option choices and the uncertainty level (*r*_RT-uncertainty_) in the initial decision was highly correlated with the uncertainty sensitivity (*A*_ROC_) across the participants (Figure 5B; *r* = 0.61, *z* test, *z* = 3.4, *P* = 4.0 × 10^−4^ in the Sudoku task; *r* = 0.48, *z* = 2.4, *P* = 0.0085 in the RDM task). Thus, the RT-uncertainty correlation also reflected individual uncertainty sensitivity.

The subjective beliefs of decision uncertainty were much reduced by redecision. The more uncertain the first decision was, the more reduced the uncertainty level was (Figure 1F). The extent of uncertainty reduction by redecision was highly correlated with the uncertainty level of the initial decision (one tailed *t* test, Goodman and Kruskal’s *γ* = 0.82 ± 0.11, *t*_21_ = 8.8, *P* = 2.1 × 10^−8^ in the Sudoku task; *γ* = 0.78 ± 0.14, *t*_21_ = 7.7, *P* = 8.2 × 10^−8^ in the RDM task). Accordingly, the objective accuracy of decisions was also improved with uncertainty reduction (Figure 1G; *r* = 0.54 ± 0.13, *t*_21_ = 4.2, *P* = 2.3 × 10^−4^ in the Sudoku task; *r* = 0.39 ± 0.14, *t*_21_ = 2.8, *P* = 5.6 × 10^−3^ in the RDM task). One may suspect that the improvement of uncertainty reduction and accuracy change would be caused by regression toward mean: the worse at the first measurement, the greater of the improvement at the second measurement. However, their decision accuracy and uncertainty levels in the final decision remained significantly differential across the different uncertainty levels of the initial decision (Figure 1C, *r* = 0.35 ± 0.15, *t*_21_ = 2.1, *P* = 0.032 in the Sudoku task; *r* = 0.36 ± 0.14, *t*_21_ = 2.6, *P* = 8.9 × 10^−3^ in the RDM task; Figure 1G, *r* = 0.32 ± 0.14, *t*_21_ = 2.0, *P* = 0.042 in the Sudoku task; *r* = 0.32 ± 0.15, *t*_21_ = 2.2, *P* = 0.028 in the RDM task), indicating that the participants’ performance in redecision reflected their (metacognition) abilities, rather than by chances. Although both uncertainty levels and decision accuracy were much improved by redecision, the divergence between the uncertainty sensitivity and the decision accuracy remained significant in the final decision (Figure 1H; one tailed paired-*t* test, *t*_21_ = 3.4, *P* = 0.0013 in the Sudoku task; *t*_21_ = 2.6, *P* = 0.0084 in the RDM task). Indeed, neither the individual uncertainty sensitivities, nor those of individual differences, were altered by redecision (Figure 1I; two tailed paired-*t* test, *t*_21_ = 0.82, *P* = 0.21 in the Sudoku task; *t*_21_ = 1.0, *P* = 0.15 in the RDM task). Similarly, neither the individual RT-uncertainty correlation coefficients, nor those of individual differences, were altered by redecision (Figure 1E; two tailed paired-*t* test, *t*_21_ = −0.77, *P* = 0.22 in the Sudoku task; *t*_21_ = 0.35, *P* = 0.36 in the RDM task). Altogether, the individual uncertainty sensitivity appeared stable and intrinsic to individual metacognition ability, independent of the decision-making process or accumulated evidence.

### The metacognition network involved in metacognitive monitoring and controlling in redecision

Commonly across the two tasks, the brain activations in the initial decision were mainly restricted to the brain areas posterior to the PFC, and the posterior part of the PFC, in particular, inferior frontal junction (IFJ) (Figure 2A, Figure S1A and S1C), while a frontoparietal control network, consisting of dACC, lFPC, anterior insular cortex (AIC), middle dorsolateral PFC (mDLPFC) and anterior inferior parietal lobule (aIPL), was newly or more extensively recruited in redecision (Figure 2B; Figure S1B and S2; Table S1). However, when a new Sudoku problem or a new RDM stimulus was presented during the redecision phase, preceded by the control conditions in the decision phase, the regions of the anterior PFC (*i.e.*, lFPC, mDLPFC, and dACC) were not activated (fMRI2, n = 17; Figure S1A and Figure S3). This result supports that the frontoparietal control network, in particular, the regions of lFPC, mDLPFC and dACC in the anterior FPC, were predominately involved in the redecision process, but not involved in the initial decision process. Thus, the redecision process evoked a separate neural system, separate from the decision-making neural system.

**Figure 2.**
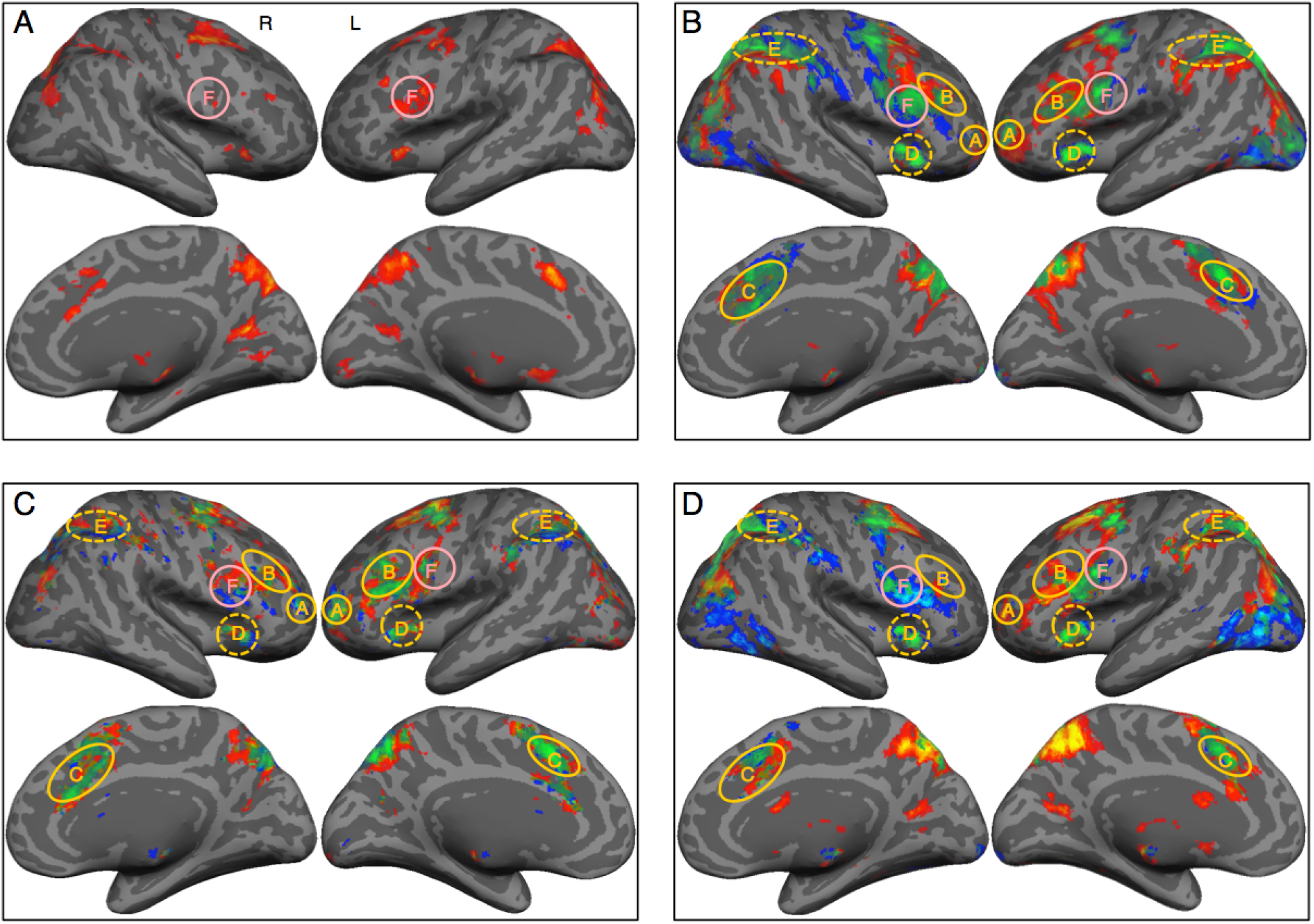
The metacognition network involving in metacognitive monitoring and metacognitive controlling. (A) Activations of the task trials in comparison with those of the control trials during the initial decision. (B) Activations of the task trials in comparison with those of the control trials during the redecision phase. (C) Activations of the task trials during the redecision phase regressed with the uncertainty levels. (D) Activations of the task trials with redecision required in comparison with those with redecision not required in fMRI3. Red-yellow patches indicate activations in the Sudoku task, blue-lightness patches indicate activations in the RDM task, *P* < 0.05, false discovery rate (FDR) corrected. Green-lightness patches indicate conjunction activations across the two tasks, *P* < 0.005, cluster-size corrected. *A*, lFPC; *B*, mDLPFC; *C*, dACC; *D*, AIC; *E*, aIPL; *F*, IFJ.

Activities in the regions of the frontoparietal control network in redecision were positively correlated with the uncertainty level of the initial decision (Figure 2C and Table S2). Critically, these correlations remained significant even for the correct trials only (Figure S1E), indicating these regions involved in uncertainty monitoring, rather than error monitoring. In contrast, activities in the ventromedial PFC (VMPFC) and posterior cingulate cortex (PCC) regions of the default-mode network were negatively correlated with the uncertainty level (Figure S1F). Although activations in the dACC and AIC regions during the decision phase were also detected by the general linear modeling (GLM) analyses (Figure 2A and Figure S1A), but activities in the two regions were not correlated with the uncertainty level (Figure S1D). Furthermore, the activations of the frontoparietal control network in redecision were not merely involved in uncertainty monitoring. In the third fMRI experiment (fMRI3, n = 25), we confirmed that the strength of activities in these regions depended critically on whether redecision on the previous situation was required after the initial decision or not. When the uncertainty levels of the initial decisions were matched in the two conditions (two tailed paired *t* test, *t*_25_ = 0.62, *P* = 0.27), activities were much stronger in the condition where redecision on the previous situation was required, in comparison with those in the condition where redecision was not required (Figure 2D), though the activities in the latter condition were also significant, and correlated with the decision uncertainty level (Figure S1G, Wan et al., 2016). Thus, the regions in the frontoparietal control network, which were more strongly activated in redecision, should be also involved in metacognitive controlling. We then putatively defined this frontoparietal control network as the *metacognition network*.

As the extent of uncertainty reduction through redecision was highly correlated with the uncertainty level of the initial decision (Figure 2F), activities in the regions of the metacognition network were also positively correlated with the extent of uncertainty reduction (Figure S1H). However, these correlations in the regions of the metacognition network became much reduced after regressed out the factor associated with the uncertainty level (Figure S1I). Conversely, the correlations with the uncertainty level remained significant after regressed out the factor associated with the extent of uncertainty reduction (Figure S1J). These partial correlation results, thus, complementarily confirmed that the cognitive processes of the regions in the metacognition network during redecision were not only comprised of metacognitive controlling, but also metacognitive monitoring. The two processes interacted with each other in redecision.

### Dissociation of metacognitive monitoring and controlling in the metacognition network in redecision

However, these two interactive processes could be dissociated in redecision. In the region that was essentially involved in uncertainty monitoring, the activity strength should dynamically reflect the extent of uncertain states. As the uncertainty was reduced by redecision, the activity strength should become weaker, and this activity change should be negatively correlated with the extent of uncertainty reduction. Alternatively, in the region that was critically involved in metacognitive controlling, the activity should become positively correlated with the extent of uncertainty reduction, representing the effort involved in metacognitive controlling. We found that the late activities in the dACC and AIC regions became negatively correlated with the extent of uncertainty reduction after orthogonalization with the uncertainty level (Figure 3A, Figure S1K and S1L). Conversely, the lFPC activity was positively correlated with the extent of uncertainty reduction after orthogonalization with the uncertainty level in the Sudoku task (Figure 3B), but negatively in the RDM task (Figure 3B and Figure S1I). These results suggest that lFPC should be instead involved in decision adjustment to reduce decision uncertainty in redecision, in particular, in the Sudoku task. In addition, the activities of the bilateral ventral IPL regions and VMPFC were also positively correlated with the level of uncertainty reduction in both tasks (Figure S1I). The VMPFC activities appeared intrinsically anti-correlated with activities of dACC or the other regions of the metacognition network (the details about the VMPFC activities will be discussed in another study). Thus, dACC and AIC appeared specifically involved in metacognitive monitoring. Instead, lFPC appeared specifically involved in metacognitive controlling. Their functional roles in metacognition were dissociated in redecision.

**Figure 3.**
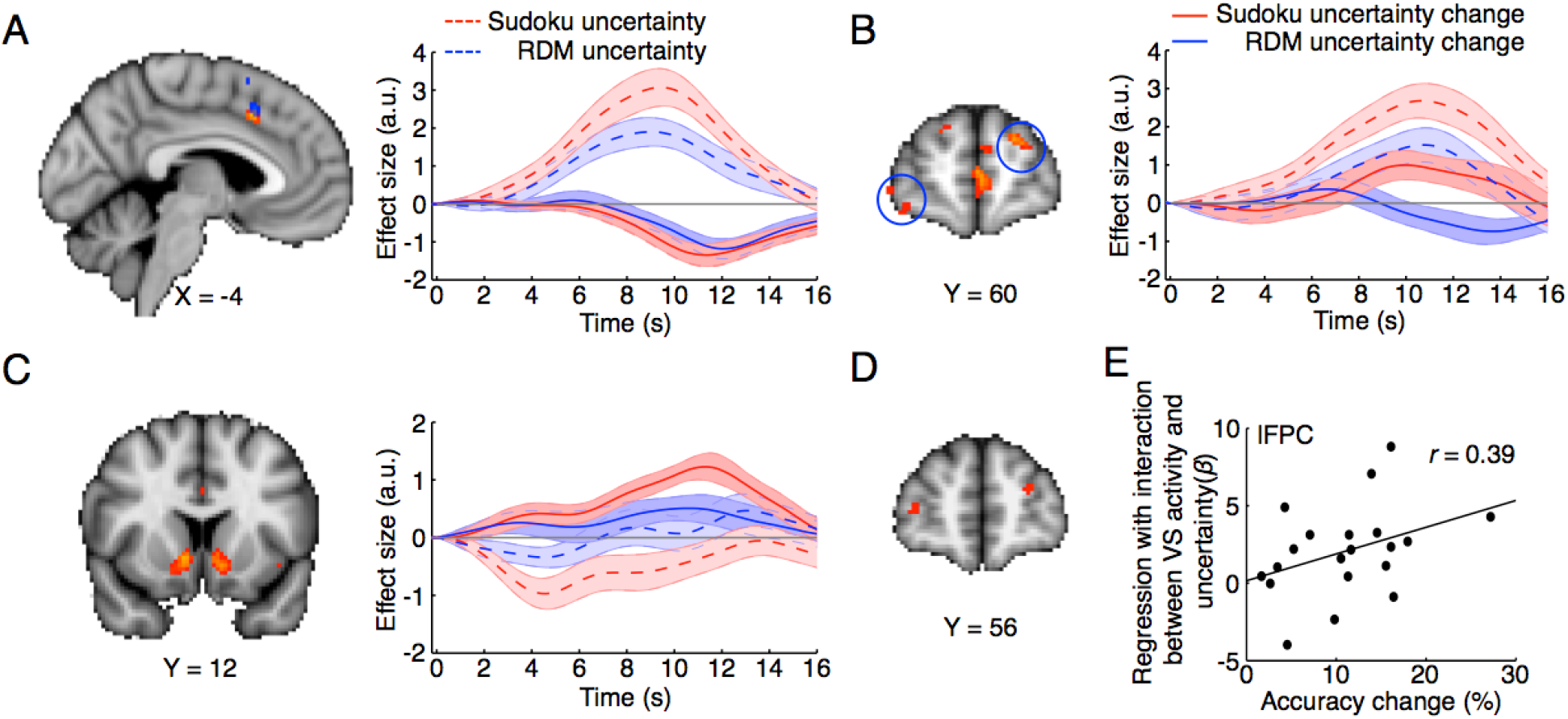
Dissociation of metacognitive monitoring in dACC and metacognitive controlling in lFPC in redecision. (A) The dACC activity was negatively correlated with the level of uncertainty reduction after orthogonalization with the uncertainty level in the Sudoku and RDM tasks. (B) The lFPC activity was positively correlated with the level of uncertainty reduction in the Sudoku task, but the correlation was negative in the RDM task. (C) The ventral striatum (VS) activity was positively correlated with the uncertainty reduction in the Sudoku task, though the early VS activity was negatively correlated with the uncertainty level. (D) The lFPC activity was significantly modulated by the VS activity (physiological effect) and the uncertainty level (psychological effect) interaction (PPI) in the Sudoku task. (E) The individual accuracy change by redecision was positively correlated with the PPI coupling strength in the lFPC region in the Sudoku task. The time courses are relative to the onset of the initial decision.

The opposite regression of the lFPC activities with the extent of uncertainty reduction in the Sudoku and RDM tasks might reflect its different roles in decision adjustment in the two tasks. Decision adjustment in the perceptual decision-making tasks would merely require low-level cognitive control, for instance, paying more attention on the new sensory information in redecision, whereas that in the rule-based decision-making tasks (i.e., Sudoku solving) would require high-level cognitive control, for instance, exploring alternative solutions. In the latter case, metacognitive controlling needed more effort, and whether the problem would be better solved should be conditioned to individual intrinsic motivation to engage the metacognitive controlling process. The ventral striatum (VS) was positively correlated with the extent of uncertainty reduction in the Sudoku task, but not in the RDM task (Figure 3C). To the end, VS might encode intrinsic motivation to engage the metacognitive controlling in the Sudoku task. Critically, the lFPC activity was significantly coupled with the interaction between the VS activity and the uncertainty level (Figure 3D; see PPI analysis in Materials and Methods), and the accuracy change of each participant by redecision was positively correlated with the coupling strength in the Sudoku task (Figure 3E). These results imply that the efficiency of lFPC involvement in metacognitive controlling in the rule-based decision-making tasks (Sudoku) should be conditioned to intrinsic motivation, modulated by the VS activity.

### Dissociation of individual metacognitive abilities of monitoring and controlling in the metacognition network

Metacognitive abilities of monitoring and controlling behaviorally embody in two components: uncertainty sensitivity and accuracy change, respectively. Through all sessions of fMRI and other repeated behavioral experiments, the individual uncertainty sensitivity was highly consistent between different sessions of the Sudoku task (Cronbach’s *α* = 0.91, Figure 4A, left column, upper panel) and the RDM task (*α* = 0.89, Figure 4A, left column, middle panel), as well as across the two tasks (*α* = 0.85, Figure 4A, left column, lower panel). In contrast, the individual accuracy change by redecision was not consistent across the two tasks (*α* = 0.03, Figure 4A, right column, lower panel), though it was consistent between different sessions of the Sudoku task (*α* = 0.80, Figure 4A, right column, upper panel) or the RDM task (*α* = 0.76, Figure 4A, right column, middle panel). Thus, individual metacognitive abilities of monitoring appeared reliably consistent, but those of metacognitive controlling were dissociated between the two tasks.

**Figure 4.**
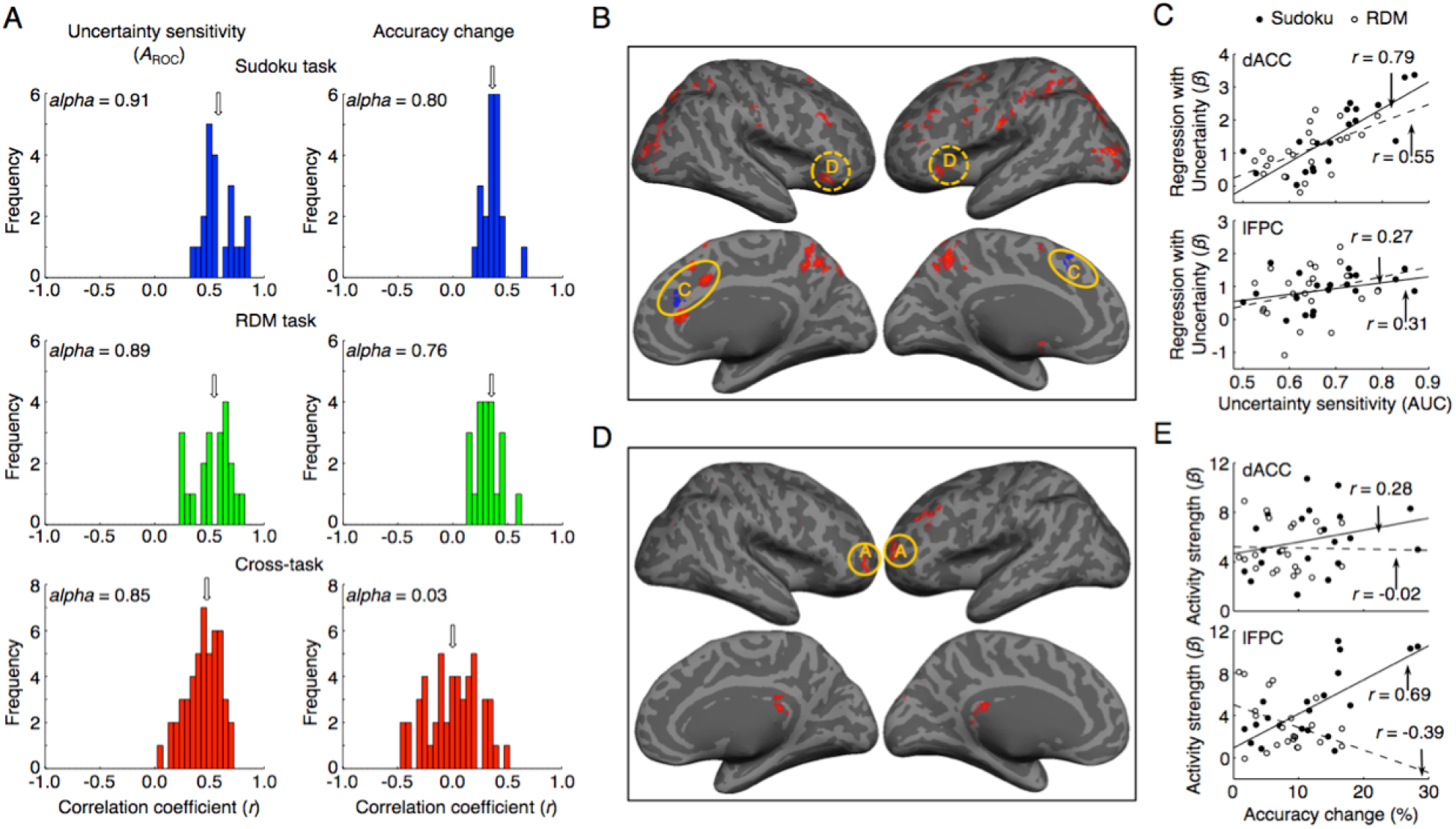
Individual metacognitive abilities of uncertainty sensitivity and accuracy change were separately associated with the dACC and lFPC activities in redecision. (A) The histograms of correlation coefficients of individual uncertainty sensitivity (*A*_ROC_, left column) and individual accuracy change (right column) between different sessions in the Sudoku or RDM task, and across the two tasks. The arrows indicate the medians of the histograms. (B) The individual uncertainty sensitivity (*A*_ROC_) was positively correlated with the uncertainty-level regression *β* values of the fMRI activities mainly in the dACC and AIC. (C) The scatter plots of the dACC and lFPC activities regressed with the uncertainty level against the individual uncertainty sensitivity. (D) The individual accuracy change was positively correlated with the mean activity predominately in the lFPC region. (E) The scatter plots of the dACC and lFPC mean activity against the individual accuracy change. In C and E, the solid lines indicate fitting data in the Sudoku task and the broken lines indicate fitting data in the RDM task. The conventions in B and C are the same as in Fig. 2.

Intriguingly, the individual uncertainty sensitivity (*A*_ROC_) was positively correlated with the uncertainty-level regression *β* value of the fMRI signal changes primarily in the dACC and AIC regions (Figure 4B and Figure 4C upper; one tailed *t*-test, *r* = 0.79, *t*_19_ = 5.6, *P* = 6.0 × 10^−6^ in the Sudoku task; *r* = 0.55, *t*_19_ = 2.9, *P* = 0.0049 in the RDM task; Table S3), but not with that in the lFPC region (Figure 4B and Figure 4C bottom; one tailed *t*-test, *r* = 0.27, *t*_19_ = 1.2, *P* = 0.12 in the Sudoku task; *r* = 0.31, *t*_19_ = 1.4, *P* = 0.085 in the RDM task), commonly in both tasks. In contrast, the individual accuracy change was significantly correlated with the mean activity in the lFPC region (Figure 4D and Figure 4E bottom; one tailed *t*-test, *r* = 0.69, *t*_19_ = 4.2, *P* = 2.2 × 10^−4^ in the Sudoku task; *r* = −0.39, *t*_19_ = 1.9, *P* = 0.041 in the RDM task), but not with that in the dACC region (Figure 4D and Figure 4E upper; one tailed t-test, *r* = 0.28, t_19_ = 1.3, *P* = 0.11 in the Sudoku task; *r* = −0.02, t_19_ = 0.09, *P* = 0.47 in the RDM task). When the lFPC activity was stronger, the accuracy change was more in the Sudoku task, but became less in the RDM task (Figure 4E). In addition, the individual accuracy change was also positively correlated with the uncertainty-level regression *β* value of the lFPC activity in the Sudoku task (Figure S4, one tailed t-test, *r* = 0.70, *t*_19_ = 4.3, *P* = 0.00017), but not in the RDM task (Figure S4, one tailed *t*-test, *r* = −0.02, *t*_19_ = 0.09, *P* = 0.47). Thus, the dACC activity (AIC as well) commonly represented individual metacognitive abilities of monitoring, whereas lFPC differentially modulated individual metacognitive abilities of controlling in the two tasks.

The RT of option choices in the initial decision after orthogonalization with the uncertainty level remained significantly correlated with the activities of the regions in the metacognition network during the redecision phase in both tasks (Figure 5A). The *r*_RT-uncertainty_ strength in each participant was highly correlated with the individual uncertainty sensitivity (*A*_ROC_) of either the initial decision or the final decision, respectively (Figure 5B). After orthogonalization with the individual uncertainty sensitivity, the individual *r*_RT-uncertainty_ strength in the Sudoku task was also significantly correlated with the uncertainty-level regression *β* value of the dACC activity, but not that of the lFPC activity (Figure 5C and 5D). As the confidence reports *per se* are subjective, the association with RT could be more objective to reflect the internal uncertain states. Altogether, these neural correlates of individual differences in metacognitive abilities further suggest that the functional roles of dACC and lFPC in metacognition should be dissociated.

**Figure 5.**
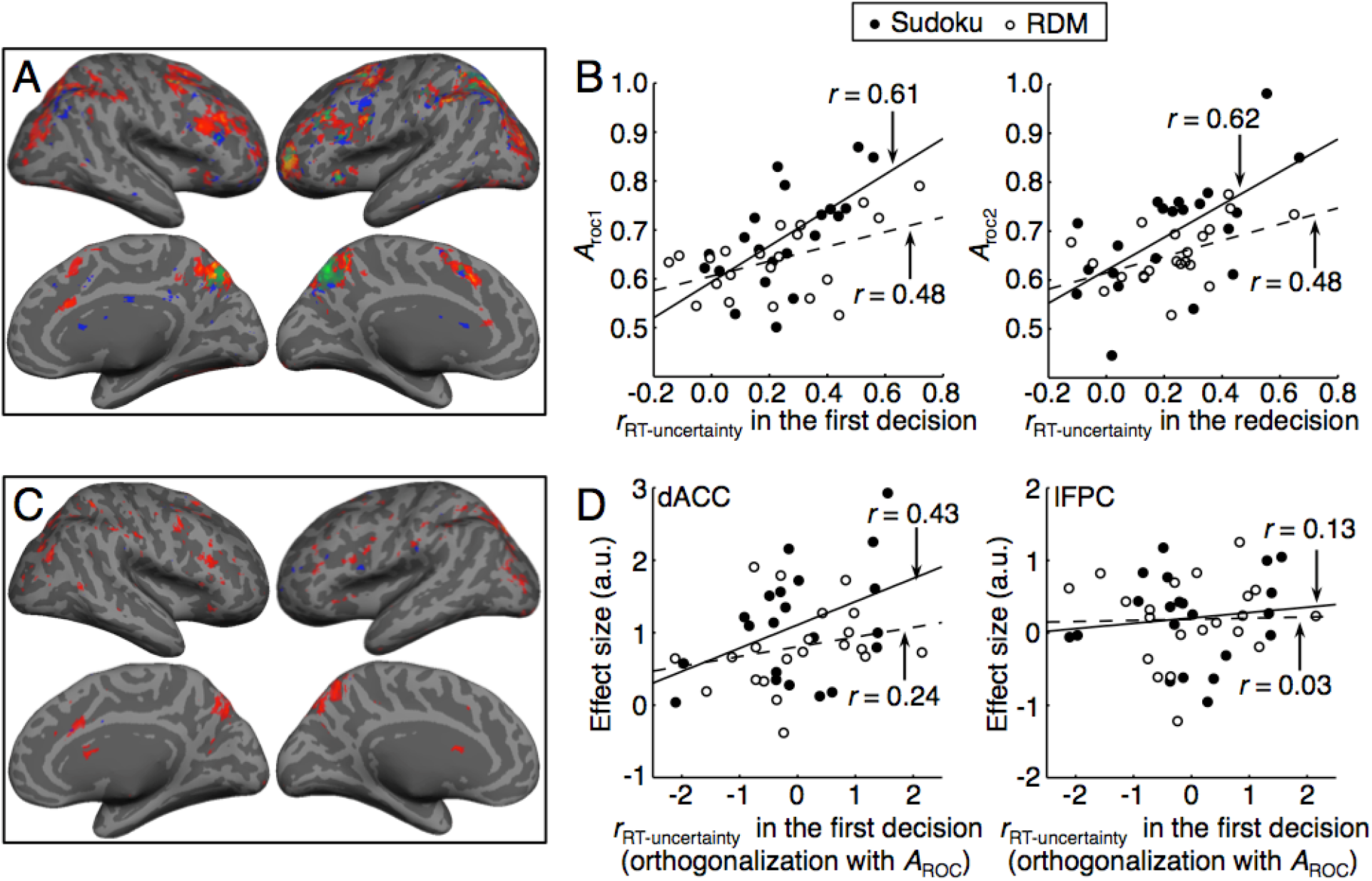
The response time (RT) also reflected decision uncertainty sensitivity. (A) The RT was positively correlated with the activities of the regions in the metacognition network in redecision, in the Sudoku and RDM tasks. (B) The individual *r*_RT-uncertainty_ was positively correlated with the individual uncertainty sensitivity (*A*_ROC_) in the initial and final decisions in both tasks. (C) The individual *r*_RT-uncertainty_, even after orthogonalization with the uncertainty sensitivity, was also positively correlated with the uncertainty-level regression *β* values of the dACC activities mainly in the Sudoku task. (D) The scatter plots of the individual *r*_RT-uncertainty_ after orthogonalization with the uncertainty sensitivity, against with the uncertainty-level regression *β* values of the fMRI activities in the dACC and lFPC regions in both tasks.

### Task baseline activities in the metacognitive network predicting individual metacognitive abilities of monitoring and controlling

The regions of the metacognition network were also activated in the certain trials of both tasks (confidence level = 4), in comparison with their respective control conditions (Figure 6B and Figure S2). These activation differences might be partially caused by different subjective uncertain states between the two conditions that were not reflected by the four-scale confidence ratings (the ceiling effect). The averaged accuracy was about 80% in the certain trials of the tasks (Figure 1C), but it was about 95% in the control conditions. However, these task baseline activities in the certain trials of the tasks also reflected the individual uncertainty monitoring bias and potential abilities of efficient metacognitive controlling. The individual uncertainty monitoring bias, as estimated by averaging the uncertainty levels of the trials in each session of the tasks, representing the individual over-confident or under-confident tendency, was consistent between different sessions in the Sudoku task (*α* = 0.95, Figure 6A, left panel), and in the RDM task (*α* = 0.94, Figure 6A, middle panel), as well as across the two tasks (*α* = 0.91, Figure 6A, right panel). The individual mean uncertainty level was positively correlated with the task baseline activity in the dACC region (Figure 6C and Figure 6F left; one tailed *t*-test, *r* = 0.50, *t*_19_ = 2.5, *P* = 0.0096 in the Sudoku task; *r* = 0.44, *t*_19_ = 2.1, *P* = 0.022 in the RDM task), but not with that in the lFPC region (Figure 6C and Figure 6F right; one tailed *t*-test, *r* = 0.18, *t*_19_ = 0.80, *P* = 0.22 in the Sudoku task; *r* = −0.04, *t*_19_ = 0.17, *P* = 0.43 in the RDM task), commonly in both tasks. Meanwhile, the individual accuracy change in the Sudoku task was positively correlated with the task baseline activity in the lFPC region (Figure 6D and Figure 6G right; one tailed *t*-test, *r* = 0.45, *t*_19_ = 2.2, *P* = 0.020), but not with that in the dACC region (Figure 6G left; one tailed *t*-test, *r* = 0.14, *t*_19_ = 0.62, *P* = 0.27). In contrast, the individual accuracy change in the RDM task was negatively correlated the task baseline activity in the lFPC region (Figure 6E and Figure 6G right; one tailed *t*-test, *r* = −0.40, *t*_19_ = 1.9, *P* = 0.035), but not with that in the dACC region (Figure 6G left; one tailed *t*-test, *r* = −0.13, *t*_19_ = 0.57, *P* = 0.29). Thus, the task baseline activity in the dACC region could reflect the individual uncertainty monitoring bias in both tasks, whereas that in the lFPC region could predict the individually differential potential abilities of metacognitive controlling for decision adjustment in both tasks.

**Figure 6.**
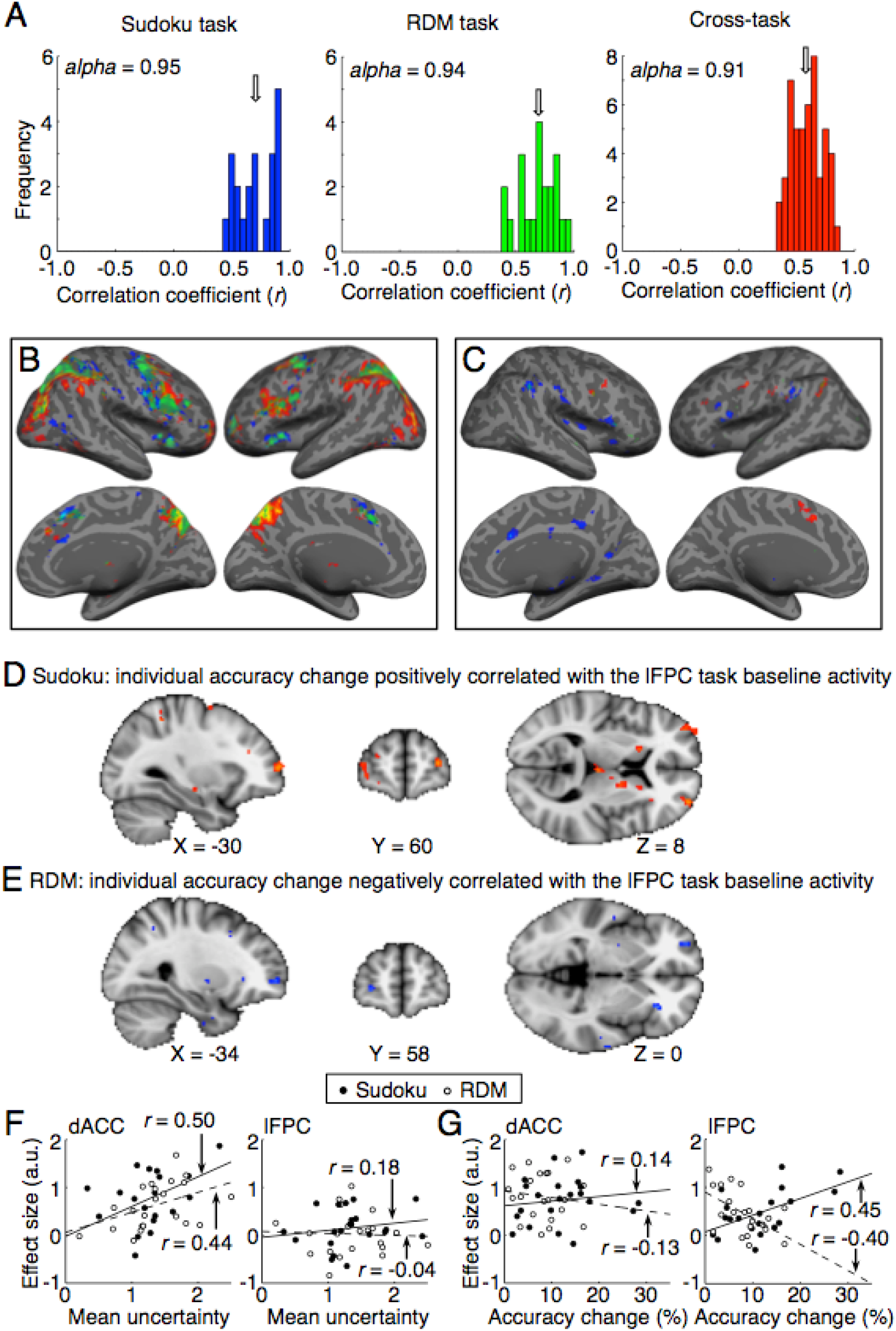
Individual metacognitive abilities predicted by the task baseline activities of the dACC and lFPC regions in the metacognition network in redecision. (A) The histograms of the correlation coefficients of individual mean uncertainty levels that represented individual bias of uncertainty sensitivity between different sessions of the Sudoku or RDM task, and across the two tasks. The arrows indicate the medians of the histograms. (B) The task baseline activities (confidence level = 4) in comparison to those of the control trials in the Sudoku and RDM tasks. (C) Positive correlation of task baseline activities during the redecision phase of the task trials with the individual mean uncertainty level across the participants. The conventions in B and C are the same as in Fig. 2. (D) The lFPC task baseline activities (in comparison to those of the control trials) were positively correlated with the individual accuracy change across the participants in the Sudoku task. (E) The lFPC task baseline activities were negatively correlated with the individual accuracy change across the participants in the RDM task. (F) The scatter plots of the dACC and lFPC task baseline activities against the individual mean uncertainty level. (G) The scatter plots of the dACC and lFPC task baseline activities against the individual accuracy change. In F and G, the solid lines indicate fitting data in the Sudoku task and the broken lines indicate fitting data in the RDM task.

### Functional connectivity in the metacognition network

Thus far we have shown that the neural system of metacognition can be dissociated into at least two subsystems: the dACC and AIC regions involved in metacognitive monitoring of decision uncertainty, and the lFPC region involved in metacognitive controlling of decision adjustment. To further elaborate the subsystems of the metacognition network, we made analyses of interregional functional connectivity in the metacognition network. By regressing out the mean activities, and the modulations by the uncertainty level, the RT and the level of uncertainty reduction, as well as their interactions, we calculated trial-by-trial correlation between each pair of regions in the metacognition network (see Materials and Methods). The interregional functional connectivity patterns in both the task condition (Figure 7A) and the control condition (Figure 7B) were almost identical between the two tasks, and also similar to that at the resting state (Figure 7C). The interregional functional connectivity patterns consistently showed that the metacognition network might be divided into three subsystems: the lFPC region; the dACC and AIC regions; the DLPFC and aIPL regions. The interregional functional connectivity within each of the subsystems was considerably stronger than that across the subsystems. So far, the functional roles of the subsystem consisting of the DLPFC and aIPL regions in metacognition remain unclear. It is worthy of noting that the functional connectivity between dACC and the regions of the other two subsystems in the task conditions was slightly stronger than the corresponding one at the resting state.

**Figure 7.**
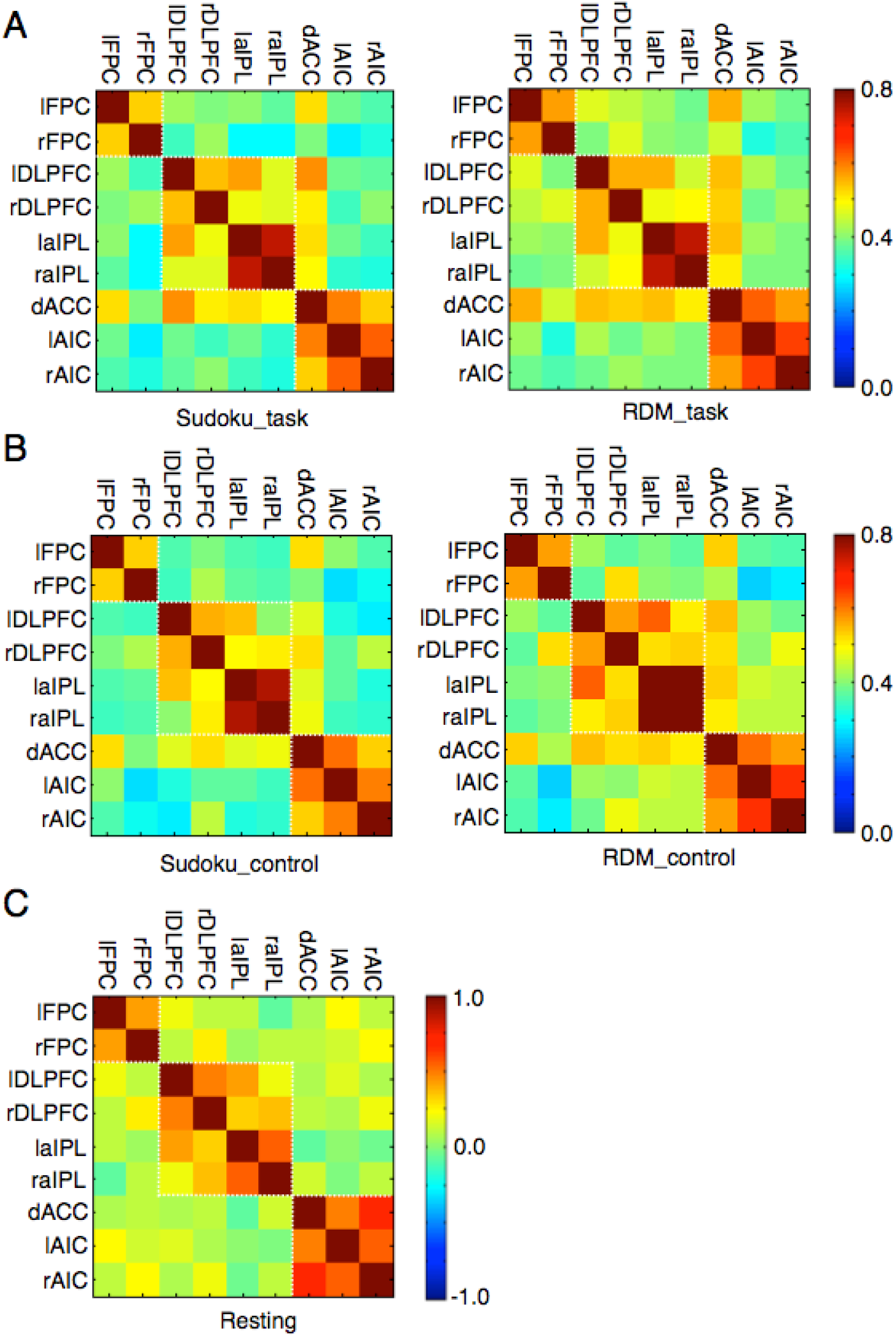
The regional functional connectivity of the metacognition network during the task (A) and control (B) conditions in the Sudoku and RDM tasks, as well as during the resting state (C).

## Discussion

In the present study, we utilized a novel “decision-redecision” paradigm to examine the behavioral and neural correlates of metacognition in decision uncertainty monitoring and decision adjustment controlling during the redecision phase, in comparison with those correlates of the decision-making process during the initial decision phase. The behavioral results were similar between the two tasks, and largely contradicted the predictions by the theory that metacognition is merely based on the very same decision-making process (Vickers 1979; Kiani and Shadlen, 2009; Resulaj et al., 2009; Pleskac and Busemeyer, 2010; Kiani et al., 2014; van den Berg et al., 2016). Given a quite longer duration for accumulating more information in redecision, the divergence between the uncertainty sensitivity and the final decision accuracy remained outstanding. Instead, our robust finding from the behavioral results was that the individual uncertainty sensitivity (both *A*_ROC_ and *r*_RT-uncertainty_) remained markedly stable between the two consecutive decisions on the same situations, between different sessions of the same tasks, and across the tasks (Song et al., 2011), indicating that the individual uncertainty sensitivity was largely independent of the accumulating evidence and the forms of the decision-making process. This leads us to favor the alternative theory that metacognition entails a separate neural system to monitor and control the decision-making neural system (Flavell, 1979; Nelson and Narens, 1990; Dunlosky and Metcalfe, 2009; Fleming and Dolan, 2012). Using fMRI, a frontoparietal control network predominately recruited in redecision was identified. This network was involved in both metacognitive monitoring of decision uncertainty and metacognitive controlling of decision adjustment, commonly in both tasks. Therefore, we putatively referred to this network as the metacognition network, which could be probably segregated into three subsystems (Figure 7).

The subsystem consisting of the dACC and AIC regions was involved in decision uncertainty monitoring, commonly in the tasks. The neural uncertainty sensitivity (the uncertainty-level regression *β* value of neural activity) in the two regions was highly correlated with the behavioral uncertainty sensitivity. Further, their task baseline activities could predict the individual uncertainty bias. Thus, the subjective uncertainty level could be represented by the dACC and AIC activities, which could transform or read out the uncertainty information from the different decision-making processes (Kepecs et al., 2008; Kiani and Shadlen, 2009; Middlebrooks and Sommer, 2012; Komura et al., 2013; Pouget et al., 2016). Although the fMRI signals of the decision-making neural system in the initial decision were not directly correlated with the uncertainty level (Pouget et al., 2016), the observation that the correlation of RT with uncertainty was significant and stable indicates that participants should be aware of decision uncertainty during choices. Together, we infer that uncertainty monitoring might be indeed consisted of two-order processes, the first-order process might coincide the decision-making process to bring out the uncertainty information (Vickers 1979; Kiani and Shadlen, 2009; Resulaj et al., 2009; Pleskac and Busemeyer, 2010; Kiani et al., 2014; van den Berg et al., 2016), and the second-order process might transform the uncertainty information from different decision-making processes into a common subjective feeling, encoding in the dACC and AIC regions. This hypothesis then integrates the two previous theories together and consistently accounts for the observed evidences from both sides. It is worthy of noting that our results were different from the previously neuroanatomical studies showing that the lFPC region was associated with the individually behavioral uncertainty sensitivity (Fleming et al., 2010; Fleming et al., 2014).

The dACC and AIC regions have been well recognized in involving conflict and error monitoring of the cognitive processes to signal the need for further control (Botvinick et al., 2001; Ridderinkhof et al, 2004; Shenhav et al., 2013). Here we demonstrated that it was decision uncertainty, rather than decision errors, to be served as the primary signal to be monitored (Wan et al., 2016). While conflict situations often but not necessarily cause uncertainty, it needs further studies to confirm whether the uncertainty information should be also critical in conflict situations. The dACC and AIC regions are shown to broadly monitor subjective feelings of such as pains, emotions and others (Crag, 2009). Critically, the salient information to elicit conscious monitoring in these regions is not necessarily from the somatosensory stimulation (Singer et al., 2004). Similarly, the prospective monitoring of uncertainty in judgments of learning (JOL) and feeling-of-knowing (FOK) also activated these regions, prior to execution of the decision-making tasks (Maril et al., 2001). Therefore, the decision uncertainty monitoring in the dACC and AIC regions should be domain-general, commonly for different forms of decision-making tasks. In turn, the uncertainty sensitivity is a unique and core trait of each individual decision maker, dependent on the circuit of the dACC and AIC regions (Craig, 2009).

Decision uncertainty monitoring could be a bottom-up process. It automatically occurred even with no requirement for redecision (fMRI3, Figure S1G; Wan et al., 2016). However, the subsequent decision adjustment should need top-down cognitive control. The activities of lFPC, rather than dACC or AIC, were positively associated with the extent of uncertainty reduction and the accuracy change by redecision in the Sudoku task, suggesting that the lFPC subsystem should be critically involved in metacognitive controlling, in particular, in the Sudoku task. Uncertainty-driven exploration could be a critical process in metacognitive controlling (Yoshida and Ishii, 2006; Daw et al., 2006; Boorman et al., 2009; Badre et al., 2012; Wan et al., 2016). To revise the foregone decisions often needs exploration of alternatives by finding an alternative solution approach, since the same solution approach as previously used in the preceding decision would very likely lead to the same solution. Thus, strategy management could be the key function of lFPC involvement in the metacognitive controlling. This top-down strategic signal might regulate the activities in the other frontal cortical areas and the posterior parietal cortex, to execute the processes of altering the previous uncertain choice (Yoshida and Ishii, 2006; Badre et al., 2012; Wan et al., 2016), or to explore a non-default option (Daw et al., 2006; Boorman et al., 2009).

Cognitive control is in general effortful (Westbrook and Braver, 2016). Decision makers tend to avoid making decisions on the tasks that are more cognitive demanding (McGuire and Botvinick, 2010), or to choose less systematic or more own suitable strategies to make decisions (Beach and Mitchell, 1978; Mattews et al., 1980). The metacognitive controlling in redecision to better solve the Sudoku problems needed the participants’ effort to engage. Since there were no external incentives to motive them to do so in the task, their engagement in metacognitive controlling should be driven by their intrinsic motivation, or curiosity. That is, to know the truth. The VS activities seem to encode this intrinsic motivation to reduce decision uncertainty, and to facilitate lFPC engagement in metacognitive controlling. This implies that dopamine might play a critical role in the lFPC activity involving in metacognitive controlling (Westbrook and Braver, 2016). How the intrinsic motivation and external incentive interacts with metacognitive controlling remains quite intriguing, and is a very important issue in education (Morgan, 1984). Alternatively, the VS activity that was positively correlated with the extent of uncertainty reduction might represent the progress of processes to achieve the goal (i.e., problem solution) through redecision (Howe et al., 2013).

It should then be much expected that lFPC would be not involved in metacognitive controlling in the RDM task, as revising the preceding perceptual decision may need no more than attention on the stimuli in redecision to accumulate new information, rather than exploration of alternative options. However, the lFPC activity remained activated too, and was negatively correlated with extent of uncertainty reduction and the accuracy change. This implies that the process of exploration in lFPC might be competitive with the simultaneous process of exploitation in the posterior brain areas when these two-level systems were not coordinated (Daw et al., 2005). Indeed, the FPC lesion on non-human primates enhanced the animals’ performance of a well-learned decision-making task (Mansouri et al., 2015). However, it remains enigmatic that lFPC was kept activated when it was not necessary and would not facilitate the engaging task. Presumably, the dACC control signals driven by decision uncertainty might non-selectively activate lFPC. The automaticity of eliciting lFPC involvement in metacognitive controlling may enhance uncertainty resolution in majority of difficult real-world situations, to relieve effort for engagement in metacognitive controlling, but failure of disentanglement however could impair the performance adjustment in simple tasks.

The Metacognitive controlling is a form of cognitive control, but not all forms of cognitive control are metacognitive. Although the Sudoku and RDM tasks appeared very different, to our surprise, the fMRI activation patterns associated with the decision-making process were quite similar between the two tasks. Critically, IFJ at the posterior PFC was commonly activated. IFJ is ubiquitously engaged in online task execution, involved in cognitive control (Brass et al., 2005; Duncan, 2010) and attention (Baldauf and Desimone, 2014). Thus, IFJ might play a critical role of object-level cognitive control generally in different decision-making tasks (Heekeren et al, 2006; Ho et al., 2009; Wan et al., 2016). The separation of the meta-level cognitive control in the anterior PFC and the object-level cognitive control in the posterior PFC is aligned with the hypothesis of the rostrocaudal functional division in the PFC (Koechlin and Summerfield, 2007; Badre and D’Esposito, 2007; Wan et al., 2016).

There were some potential pitfalls for the fMRI data analyses in the current study. As the metacognition process should automatically accompany the decision-making process with uncertainty, it excludes the conventional techniques of fMRI paradigms to insert time jitters of blank between the initial decision phase and the redecision phase. Thus, generally speaking, the two events of the decision-making process and the metacognition process in the general linear models (GLM) could be collinear, and result in inflations of standard errors of the estimated parameters, in particular, for the regions to be involved in both processes. Fortunately, for the regions of interest involved in metacognition, consistent with our predictions, their activations predominately appeared in redecision. Actually, the variance inflation factor (VIF) was about 2.4, suggesting the collinearity of the GLM models was not severe.

In summary, decision-making is usually accompanied by uncertainty. The subsequent decision uncertainty monitoring and decision adjustment tend to be automatically elicited by uncertainty. Thus, decision-making might be usually accompanied by metacognition, and the two processes are sequentially coupled together. However, the neural system of metacognition remains largely unclear so far, and was often misattributed to the decision-making process. For the first time, to the best of our knowledge, we here constructed the extent and generality of the functional architecture of the metacognition neural system in the PFC, separate from the decision-making neural system (Figure 8). The metacognition neural system is comprised of the metacognitive monitoring system and the metacognitive controlling system. The metacognitive monitoring system consisting of the dACC and AIC regions are domain-general. It reads out the uncertainty information from the decision-making process and quantitatively encodes the subjective uncertainty states. The metacognitive controlling system of the lFPC region implements high-level cognitive control (e.g., strategy), dominantly in the rule-based and abstract inference tasks (e.g., the Sudoku task), and might compete with low-level cognitive control (e.g., attention), dominantly in the perceptual tasks (e.g., the RDM task). The high-level cognitive control by the lFPC region is modulated by intrinsically motivational signals from the VS region. These two subsystems sequentially monitor and control the decision-making system, which is presumingly controlled by the IFJ region. The functions of the third subsystem of the DLPFC and aIPL regions remain to be explored in the future. Thus, the decision-making neural system and the metacognition neural system construct a closed-loop system to control and adapt our behaviors toward the task goals. Finally, Further deepening our understanding of the metacognition neural system will facilitate us to optimize the strategies for individual efficient learning and decision-making (Koriat, 1997), and help us reveal causes of metacognitive disorders in neuropsychiatric diseases (David, 1990; Dunlosky and Metcalfe, 2009).

**Figure 8.**
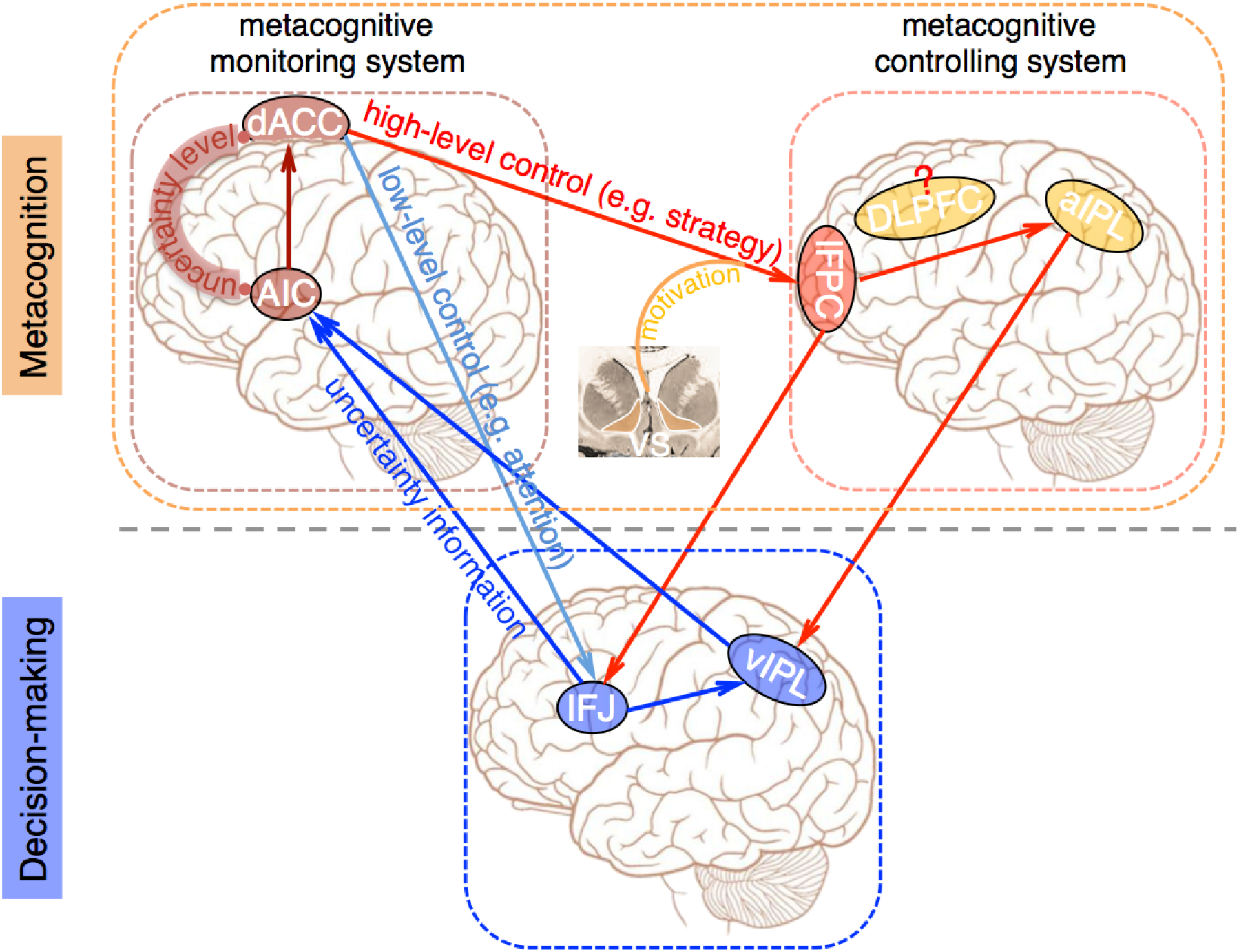
The functional architecture of the metacognition neural system. The scheme of functional architecture of the metacognition neural system and its interactions with the decision-making neural system, synthesized from the converging results in the current study. The metacognition neural system is comprised of the metacognitive monitoring system (dACC and AIC) and the metacognitive controlling system (lFPC). The decision-making neural system and the metacognition system construct a closed-loop system to control and adapt our behaviors toward the task goals.

## Materials and Methods

### Participants

All participants were university students, who were recruited through campus bulletin board system (BBS). Informed consent was obtained from each individual participant in accordance with a protocol approved by Beijing Normal University Research Ethics Committee. 21 participants (19–33 years old, 12 female) took part in the main fMRI experiment (fMRI1) and the resting fMRI experiment. Out of them, 16 participants (19–33 years old, 9 female) took part in all sessions of the repeated behavioral experiments. In addition, 17 participants (19–25 years old, 10 female) took part in the second fMRI experiment (fMRI2), and 25 participants (19–27 yeas old, 14 female) took part in the third fMRI experiment (fMRI3).

### RDM task

In an aperture with the radius of three degrees (visual angle), hundreds of white dots (radius: 0.08 degrees, density: 2.0%) were moving toward different directions with a speed of 8.0 degrees/second under a black background. The lifetime of each dot lasted for three frames. A part of dots were moving toward the same direction (one of the four directions: Left, Down, Right and Up), but the others were moving toward different random directions. The participant was required to discriminate the net motion direction. According to the proportion of coherently moving dots, the discrimination difficulty was classified into ten levels (Figure 1B), of which the coherences varied from 1.6% to 51.2%, whereas the coherence of moving dots in the control condition was 100%.

### Sudoku Task

In a 4 × 4 grid matrix, each digital number from 1 to 4 should be filled once and only once in each column, each row, and each corner with four grids. The task used in the present study was to fill in a target grid with a digital number from 1 to 4 in a partially completed Sudoku problem. Each problem had a unique solution. A Sudoku generator (custom codes) created thousands of different Sudoku problems. According to the minimum numbers of logic operation steps to arrive at the solutions, the problem difficulties were classified into ten levels, which largely matched with the participants’ subjective difficulty levels (Figure 1B). In the control condition, the presented problem was comprised of symbols (‘#’) in replace of the digital numbers other than that in the target grid where the digital number was illustrated. Thus, the participant only needed to press the corresponding button.

### Learning procedure

The participant learned the cognitive skills to solve the 4 × 4 Sudoku problems under the experimenters’ guidance for at least two hours per day in continuous four days. The participant first practiced to solve problems with free time in 2–4 runs, each of which comprised 40 problems at a certain difficulty level. Once the average accuracy of that session crossed over 90%, he/she then practiced to solve the problems at the same level in 2 s. Once the average accuracy of the run was over 70% in the time-limited task, the participant then repeated the above procedure with a task difficulty level upgraded. After four-day intensive training, each participant attained a high-level proficiency to solve the 4 × 4 Sudoku problems in 2 s, as the mean task difficulty finally approached about the fifth level.

### Task sequences

The sequences of both Sudoku and RDM tasks were identical. In fMRI1, each trial started with a green cross cue to indicate that the task stimulus would be presented 1 s later. The stimulus was presented for 2 s, and then four options were presented and the participant made a choice in 2 s. After an option was chosen, four confidence levels from 1 (lowest) to 4 (highest) were presented and the participant reported the confidence in 2 s. The same stimulus was immediately presented again for 4 s, and then the participant selected a choice and reported the confidence level again. Each trial lasted for 15 s. The control trials were intermingled with the task trials. The sequence of the control trials was identical to that of the task trials. In each task, there were 4 runs and each run consisted of 30 task trials and 10 control trials. The task difficulty of each trial was adjusted by a staircase procedure through which one level was upgraded after two consecutive correct trials and one level was downgraded after two consecutive erroneous trials, and kept as the same otherwise, so that the mean accuracy was converged to about 50%. Prior to each experiment, two runs were carried out for each participant to practice and to stabilize performance. The Sudoku problems used in the learning and practice sessions were different from those used in the fMRI and behavioral experiments. In addition, a ten-minute resting fMRI experiment was conducted when the participant was in a resting state with eyes opened.

The second fMRI experiment (fMRI2, Figure 3 and Figure S1C) was carried out to examine whether the metacognition network would be also essentially involved in the cognitive processes of the initial decision, when a new Sudoku problem or RDM stimulus was presented for decision at the first time during the redecision phase, following the control conditions in the decision phase. In the decision phase, all situations were those as used in the control conditions of fMRI1. In the redecision phase, the same control situations appeared in a half of trials and new Sudoku problems (or RDM) stimuli appeared in the other half of trials. These two cases appeared randomly in the redecision phase. The new Sudoku problems (or RDM) stimuli used in the experiment were selected from those in which each individual participant would mostly make confirmative choices, that is, the confidence ratings were predominately 4. The task sequence was same as used in fMRI1. In total, there were 120 trials across two runs.

The third fMRI experiment (fMRI3, Figure 2D and Figure S1E) was carried out to compare brain activities in the redecision condition (required to make a decision on the foregone situation again) with those in the non-redecision condition (not required to make a decision on the foregone situation again) following the initial decisions in both Sudoku and RDM tasks. The task sequence was very similar as used in fMRI1, but the presentation time of the stimulus was 3 s during the redecision phase. The stimuli used in the non-redecision condition during the second phase were those used in the control condition in each task. In each task, both the redecision and non-redecision conditions were randomly intermingled, and each consisted of 60 trials across 3 runs.

In the fMRI experiments, the participants viewed images of the stimuli on a rear-projection screen through a mirror (resolution, 1024 × 768 pixels; refresh rate, 60 Hz). Normal or corrected-to-normal vision was achieved for each participant. All images were restricted to 3 degrees surrounding the fixation cross.

### fMRI experiments

All fMRI experiments were conducted using a 3 T Siemens Trio MRI system with a 12-channel head coil (Siemens, Germany) after the four-day Sudoku training. Functional images were acquired with a single shot gradient echo T_2_* echo-planar imaging (EPI) sequence with volume repetition time (TR) of 2 s, echo time (TE) of 30 ms, slice thickness of 3.0 mm and in-plane resolution of 3.0 × 3.0 mm^2^ (field of view [FOV]: 19.2 × 19.2 cm^2^; flip angle [FA]: 90 degrees). Thirty-eight axial slices were taken, with interleaved acquisition, parallel to the anterior commissure-posterior commissure (AC-PC) line.

### Behavioral experiments

To test the reliability of the participants’ metacognitive abilities, behavioral experiments were carried out using same paradigms of the Sudoku and RDM tasks. Each of the participants repeatedly participated 6 sessions of the behavioral experiments in different days. Each session was comprised of 4 runs of the Sudoku task and 4 runs of the RDM task, as same as those of fMRI1.

### Behavioral data analyses

A nonparametric approach was employed to assess each participant’s uncertainty sensitivity. The receiver operating characteristic (ROC) curve was constructed by characterizing the incorrect probabilities under different uncertainty levels of the first decisions. The area under curve (AUC) was calculated to represent how well the participant was sensitive to their decision uncertainty (Fleming et al., 2010). The individual uncertainty bias was estimated by the mean uncertainty level of each session, regressed out the factor of *A*_roc_. The accuracy change was the change of mean accuracy from the first decision to the second decision. The individual uncertainty sensitivity and uncertainty bias, as well as accuracy change, were calculated for each session of the fMRI and behaviroal experiments.

### fMRI analyses

The analysis was conducted with FMRIB’s Software Library (FSL, Smith et al., 2004). To correct for the rigid head motion, all EPI images were realigned to the first volume of the first scan. Data sets in which the translation motions were larger than 2.0 mm or the rotation motions were larger than 1.0 degree were discarded. It turned out that no data discarded in the fMRI experiments. The EPI images were first aligned to individual high-resolution structural images, and were then transformed to the Montreal Neurological Institute (MNI) space by using affine registration with 6 degrees of freedom and resampling the data with a resolution of 2 × 2 × 2 mm^3^. A spatial smoothing with a 4-mm Gaussian kernel (full width at half-maximum) and a high-pass temporal filtering with a cutoff of 0.005 Hz were applied to all fMRI data.

Each trial was modeled with three regressors: the first regressor representing the first decision was time-locked to the onset of the first stimuli presentation with summation of the presentation time (2 s) and the differential RT from the mean RT of control trials as the event duration; the second regressor representing the second decision (redecision) was time-locked to the onset of the first confidence judgment, with summation of the confidence report, the second presentation time (4 s) of the stimuli and the differential RT from the mean RT of control trials as the event duration; the third regressor representing the baseline during the inter-trial intervals (ITI) was time-locked to the onset of ITI with the ITI duration as the event duration. The uncertainty level, the RT and the level of uncertainty reduction (differences of the uncertainty level between the final decision and the initial decision) were implemented as modulators of the second regressor (redecision) by demeaning the variances of the uncertainty level (Figure 2C) and consequently orthogonalizing the RT and the level of uncertainty reduction with each other (Figure 2A–C and Figure S1I), or reversing the orthogonalization order (Figure S1J).

For group level analysis, we used FMRIB’s local analysis of mixed effects (FLAME), which model both “fixed effects” of within-participant variance and “random effects” of between-participant variance using Gaussian random-field theory. Statistical parametric maps were generated by a threshold with *P* < 0.05 with false discovery rate (FDR) correction, unless noted otherwise. The regressions of the individual uncertainty sensitivity (*A*_ROC_), the individual RT-uncertainty correlation coefficient, the individual mean uncertainty level and the individual accuracy change with the *β* weights of uncertainty levels (Figure 4B, Figure 4D, and Figure S5C), or with the task baseline activities (Figure 5C–E), were calculated at the third-level of group analyses. For these analyses, Statistical parametric maps were generated by a threshold with *P* < 0.005 with the cluster-size threshold as 20.

### ROI analyses

The region-of-interest (ROIs) of the metacognition network were defined by the voxels that were significantly activated during the redecision phase in the task trials compared to those during the same phase in the control trials across both tasks using conjunction analysis (*P* < 0.005, cluster-wise correction; green areas in statistical parametric maps). ROI analyses were obtained from both hemispheres of the same region. The ventral striatum (VS) ROI was anatomically defined by the striatum atlas of FSL templates (Patenaude et al., 2011). The time courses were derived from the ROIs, calculating a mean time course within a ROI in each participant individually. We then averaged the time courses of the same condition across the participants (Figure S2 and Figure S3), or oversampled the time course by 10 and created epochs from the beginning of an event onward and applied a GLM to every pseudo-sampled time point separately. By averaging the *β* weights across participants we created the time courses shown in Figure 3. Standard errors of mean (S.E.M.) were calculated between participants.

### PPI analysis

The physiology-psychological interaction (PPI) analysis (Figure 3D) was conduced with the demeaned VS time courses after removing the mean activity and the component correlated with the uncertainty level as the physiological factor, and the uncertainty level convolved with the canonical hemodynamic response function (HRF) during the redecision phase as the psychological factor. The two factors *per se* and the interaction between the two factors, as confound regressors, were put together into a new GLM analysis across the whole brain.

### Functional connectivity analyses

Functional connectivity analyses were independently conducted for the task and resting fMRI data. For the task fMRI data, of each ROI, the residual time courses after regressed out the mean activity and the components associated with the uncertainty level, the RT, the level of uncertainty reduction and their interactions, were averaged across the voxels of the region and segmented into the individual trials of the task and control conditions in the Sudoku and RDM task, respectively. The segmented data of each trial were then modeled using a single regressor during the redecision phase convolved with the canonical HRF and then a regression value was obtained for each trial. The correlation coefficient of the regression values between each pair of the ROIs in the metacognition network was calculated across the trials of the task or control condition in each participant. Finally, the averaged correlation coefficients were shown (Figure 7A and 7B). For the resting fMRI data, the standard processing was carried out (Fox et al., 2005), and the averaged correlation coefficients were shown (Figure 7C).

## Authors contributions

L.Q., Y.N. and J.S. conducted the experiments; J.S. and X.W. analyzed the fMRI data; J.S. and X.W. designed the experiments; X.W. wrote the manuscript. X.W. and X.L. supervised the project.

## Acknowledgements

We thank Wenbin Jia, Xuesong Zhang, Sidong Wang for technical assistance and Dr. Kang Cheng for comments on the manuscript. This research was funded by the National Natural Science Foundation of China (No. 31471068 to X.W.; No. 61273063 to X.L.), Key Program for International S&T Cooperation Projects of China (MOST, 2016YFE0129100, X.W.), partially supported by “the Fundamental Research Funds for the Central Universities (2017EYT33, X.W.), and the Thousand Talents Program for Distinguished Young Scholars (X.W.).

---

**Figure Supplementary 1 (related to Figure 2).**
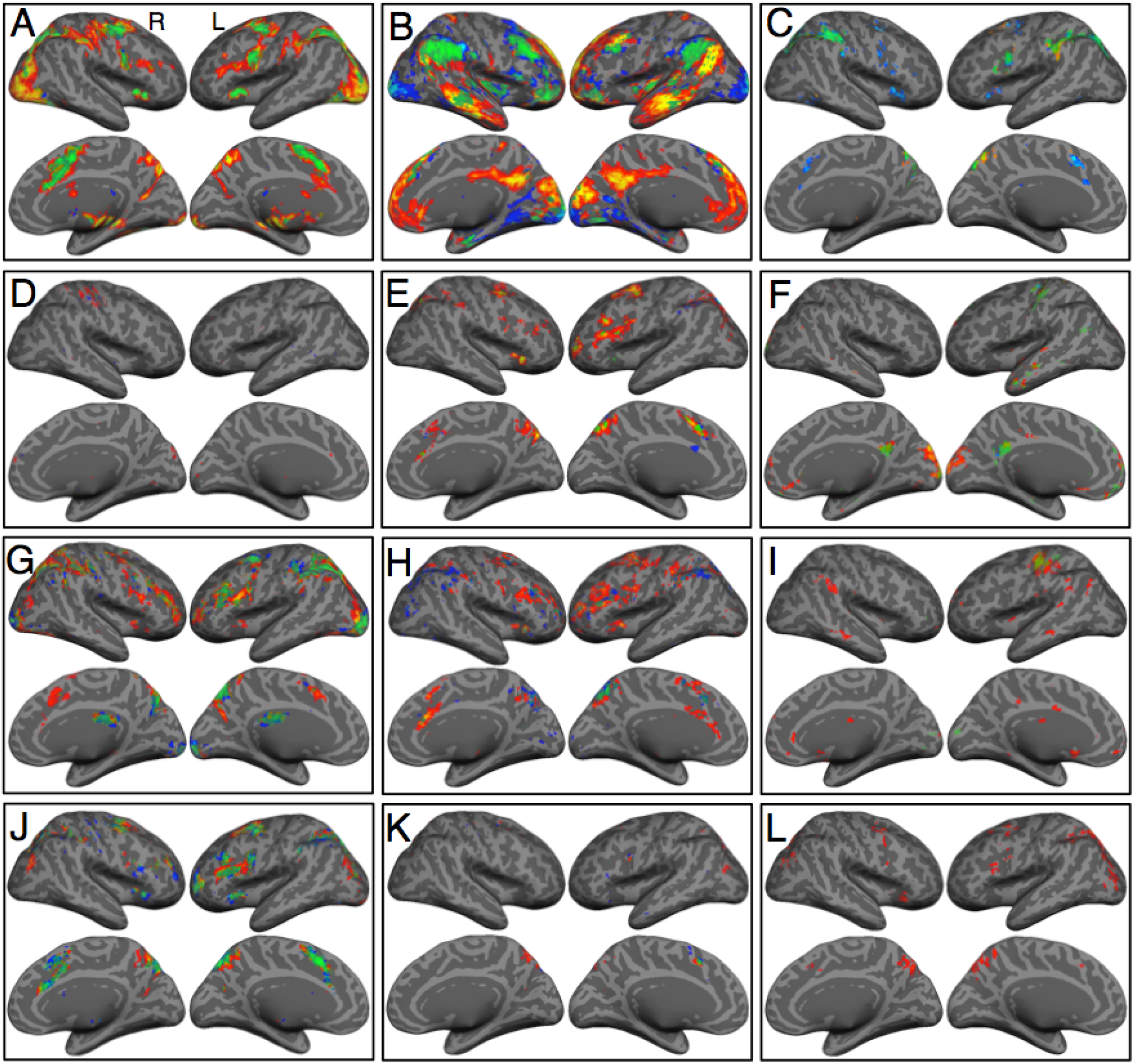
Collective statistical parametric maps in the experiments. (A) Activations during the decision phase compared with those during the ITI period in fMRI1. (B) Activations during the redecision phase compared with those during the decision phase in fMRI1. (C) Activations of the initial decision during the redecision phase compared with those of the control condition during the same phase in fMRI2. (D) Positive correlation of activities during the decision phase with the uncertainty level in fMRI1 (there were also no negative correlation). (E) Positive correlation of activities during the redecision phase of the correct trials with the uncertainty level in fMRI1. (F) Negative correlation of activities during the redecision phase with the uncertainty level in fMRI1. (G) Activations during the redecision phase without requirement to decide the previous situation again compared with those of the control trials during the same phase in fMRI3. (H) Positive correlation of activities during the redecision phase with the level of uncertainty reduction in fMRI1. (I) Positive correlation of activities during the redecision phase with the level of uncertainty reduction after orthogonalization with the uncertainty level in fMRI1. (J) Positive correlation of activities during the redecision phase with the uncertainty level after orthogonalization with the level of uncertainty reduction in fMRI1. (K) Negative correlation of activities during the redecision phase with the level of uncertainty reduction after orthogonalization with the uncertainty level in fMRI1. (L) Positive correlation of activities during the redecision phase with the interaction between the uncertainty level and the level of uncertainty reduction in fMRI1. The conventions are the same as in Fig. 2.

**Figure Supplementary 2 (related to Figure 2).**
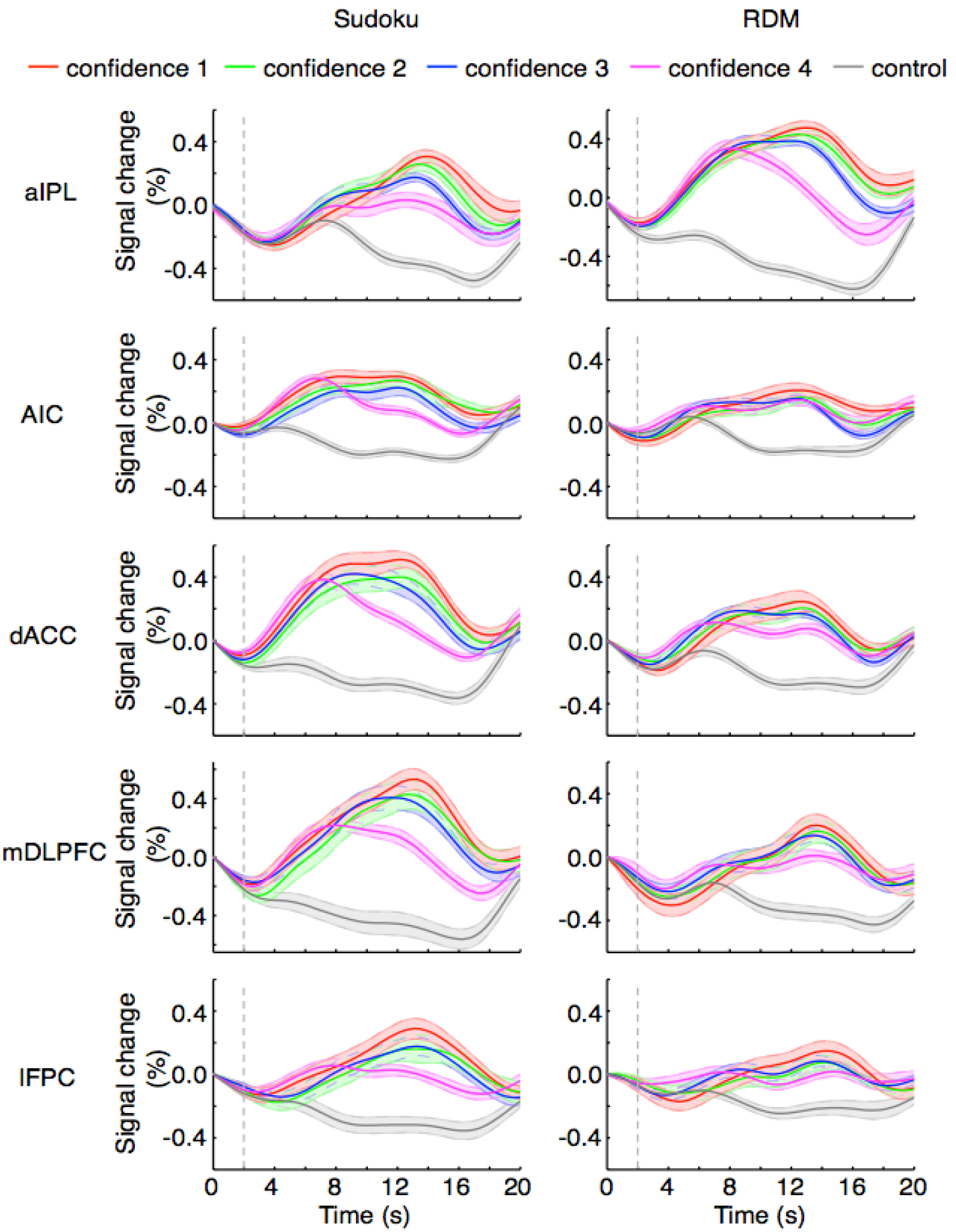
The time courses of fMRI signal changes of the regions in the metacognition network at different confidence levels in fMRI1. The time zero was the onset of the initial decision and the dash line indicates the mean offset of the initial decision.

**Figure Supplementary 3 (related to Figure 2).**
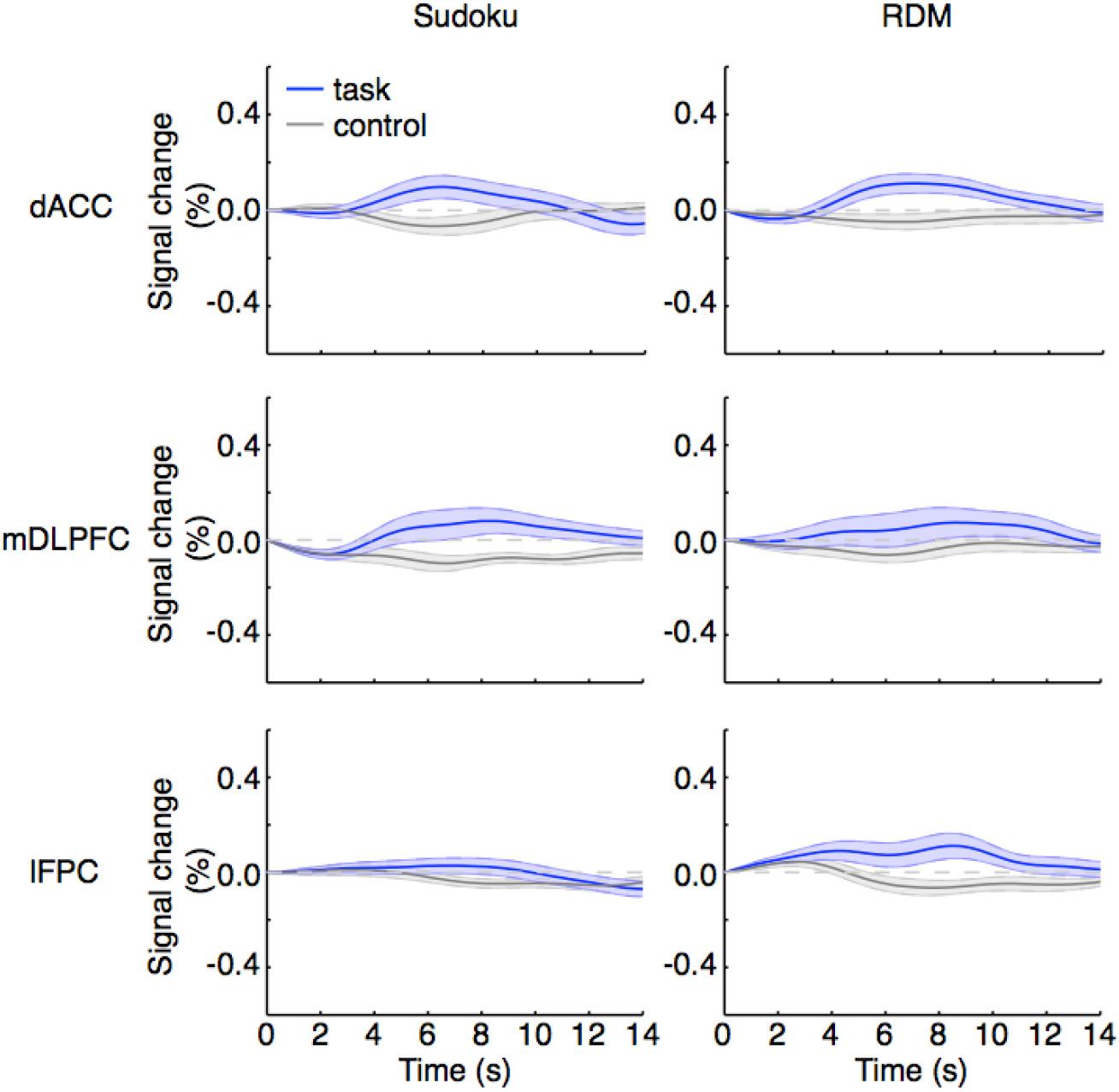
The time courses of the fMRI signal changes of the dACC, mDLPFC and lFPC regions during the initial decision in the second phase in fMRI2. The time zero was the onset of the stimuli presentation in the second phase. The participant made the initial decision in the second phase and the decision duration lasted for 4 s, longer than the initial decision period (2 s) in fMRI1. It should be noted that there were no significant activities in the mDLPFC and lFPC, whereas the dACC activities were delayed for over 3 s from the onset.

**Figure Supplementary 4 (related to Figure 4).**
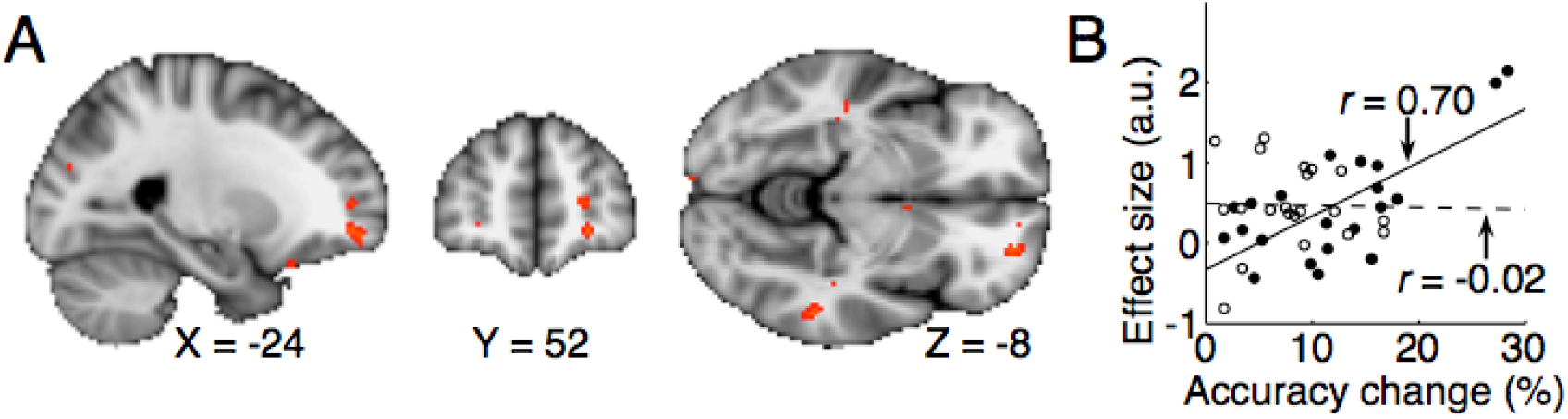
The individual accuracy change by redecision was positively correlated with the uncertainty-level regression *β* value of the lFPC activity in the Sudoku task (A; one tailed *t*-test, *r* = 0.70, *t*_19_ = 4.3, *P* = 0.00017), but not in the RDM task (B; one tailed *t*-test, *r* = 0.70, *t*_19_ = 4.3, *P* = 0.00017).

**Table S1.**
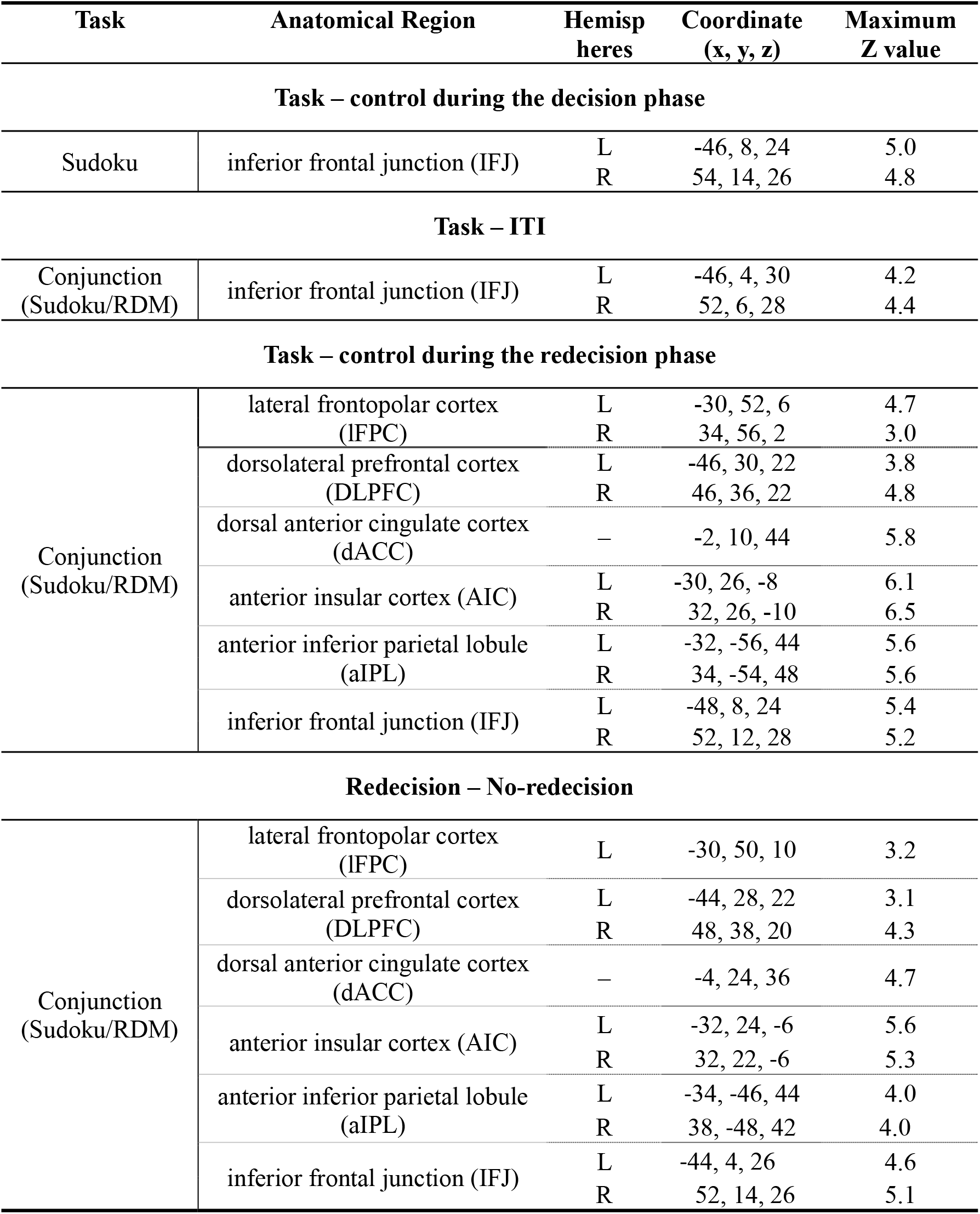
Activations between the task and control conditions. *Related to Figure 2*

**Table S2.** Activations correlated with the uncertainty level and the uncertainty reduction during the redecision phase. *Related to Figure 2–3*

| Task | Anatomical Region | Hemispheres | Coordinate (x, y, z) | Maximum |
| --- | --- | --- | --- | --- |
| **Uncertainty (positive)** |  |  |  |  |
| Conjunction<br>(Sudoku/RDM) | lateral frontopolar cortex (IFPC) | L | -30, 56, 4 | 4.2 |
|  |  | R | 30, 52, 10 | 3.5 |
|  | dorsolateral prefrontal cortex (DLPFC) | L | -44, 28, 24 | 4.4 |
|  |  | R | 42, 30, 22 | 3.7 |
|  | dorsal anterior cingulate cortex (dACC) | — | -4, 14, 46 | 4.8 |
|  | anterior insular cortex (AIC) | L | -30, 26, -4 | 4.3 |
|  |  | R | 32, 24, -2 | 4.5 |
|  | anterior inferior parietal lobule (aIPL) | L | -48, 12, 26 | 4.0 |
|  |  | R | 48, 14, 24 | 3.7 |
|  | anterior inferior parietal lobule (aIPL) | L | -32, -56, 40 | 3.9 |
|  |  | R | 44, -42, 54 | 3.8 |
| **Uncertainty (negative)** |  |  |  |  |
| Conjunction<br>(Sudoku/RDM) | ventromedial prefrontal cortex (VMFPC) | — | 0, 48, -14 | 3.8 |
|  | posterior cingulate cortex (PCC) | — | 0, -48, 22 | 4.0 |
| **Uncertainty reduction (positive)** |  |  |  |  |
| Sudoku | ventral striatum (VS) | L | -10, 12, -8 | 4.2 |
|  |  | R | 10, 12, -4 | 4.5 |
|  | ventromedial prefrontal cortex (VMFPC) | — | -2, 56, -4 | 4.2 |

**Table S3.** Activations positively correlated with the individual uncertainty sensitivity and the individual accuracy change. *Related to Figure 4–6*

| Task | Anatomical Region | Hemispheres | Coordinate (x, y, z) | Maximum |
| --- | --- | --- | --- | --- |
| **Uncertainty sensitivity ($A_{roc}$)** |  |  |  |  |
| Sudoku | dorsal anterior cingulate cortex (dACC) | R | 8, 16, 38 | 3.5 |
|  | anterior insular cortex (AIC) | L | -38, 22, -4 | 3.8 |
|  |  | R | 42, 24, -12 | 4.1 |
| RDM | dorsal anterior cingulate cortex (dACC) | L | -4, 18, 48 | 3.4 |
| **RT-uncertainty correlation coefficient<sup>1</sup>** |  |  |  |  |
| Sudoku | dorsal anterior cingulate cortex (dACC) | R | 6, 18, 42 | 3.1 |
| **Mean uncertainty** |  |  |  |  |
| Sudoku | dorsal anterior cingulate cortex (dACC) | L | -6, 16, 44 | 3.3 |
| RDM | dorsal anterior cingulate cortex (dACC) | R | 4, 14, 46 | 3.6 |
|  | anterior insular cortex (AIC) | R | 40, 22, -14 | 3.5 |
| **Accuracy change** |  |  |  |  |
| Sudoku | lateral frontopolar cortex (IFPC) | L | -26, 54, 10 | 4.0 |
|  |  | R | 26, 48, 6 | 3.7 |

<sup>1</sup>after orthogonalization with the uncertainty sensitivity.

